# Nutritional-Metabolic Lipid Profiling with LipidOne for plasma lipidomics interpretation in metabolic health

**DOI:** 10.64898/2026.05.14.725104

**Authors:** Dorotea Frongia Mancini, Husam B.R. Alabed, Roberto Maria Pellegrino

**Affiliations:** Department of Chemistry, Biology and Biotechnology, University of Perugia, Italy; Institute of Metabolic Science-Metabolic Research Laboratories, University of Cambridge, Cambridge, UK

**Author notes:** These authors contributed equally to this work (Dorotea Frongia Mancini, Husam B.R. Alabed).

## Abstract

**Background/Objectives:** Human plasma lipidomics provides valuable information on dietary and metabolic phenotypes, but the interpretation of high-dimensional lipid datasets remains challenging. We developed the Nutritional-Metabolic Lipid Profile (NMLP) module within LipidOne to translate plasma lipidomics data into interpretable nutritional-metabolic indices, functional categories, visual outputs, and biological statements.

**Subjects/Methods:** NMLP calculates lipid indices reflecting cardiometabolic lipid status, fatty acid remodelling, overall lipid quality, oxidative protection, and omega-3/essential fatty acid status. The module was applied to three human plasma lipidomics public datasets: a randomized crossover glycemic-load feeding study, a eucaloric high-fat diet intervention in normal-weight women, and a large public dataset stratified by insulin sensitivity.

**Results:** Across datasets, NMLP converted complex lipidomic matrices into coherent nutritional-metabolic profiles. In the glycemic-load study, the module highlighted metabolic lipid shifts not captured by standard clinical lipid panels, mainly involving cardiometabolic lipid status, oxidative protection, and fatty acid remodelling. In the high-fat diet intervention, NMLP tracked temporal lipid remodelling across pre-diet, on-diet, and post-diet states, consistent with metabolic adaptation to increased dietary fat exposure. In the insulin-sensitivity dataset, insulin-resistant subjects showed a storage-oriented lipid phenotype characterized by increased neutral lipid storage indices and altered lipid quality and oxidative-protection features. Category-level clustering further revealed heterogeneous nutritional-metabolic states within insulin-resistant subjects.

**Conclusions:** NMLP provides a deeper and clearer interpretative framework for human plasma lipidomics in nutrition and metabolic health research. By translating lipid species into functional indices and category-level readouts, the module may facilitate the use of lipidomics in clinical nutrition, metabolic phenotyping, and precision nutrition studies. NMLP is freely accessible as part of the online LipidOne platform.

## 1. Introduction

Human plasma lipidomics is increasingly used in nutrition and metabolic health research because circulating lipids provide a sensitive readout of dietary exposure, metabolic adaptation, and cardiometabolic risk [1]. Compared with conventional clinical lipid panels, mass spectrometry-based lipidomics semi-quantifies hundreds of lipid species across glycerolipids, glycerophospholipids, sphingolipids, sterol lipids, and related classes, offering a more detailed representation of lipid metabolism. Importantly, the field is also moving toward greater analytical standardization and clinical translation, as illustrated by recent inter-laboratory efforts showing concordant absolute quantification of clinically used plasma ceramides in reference materials and emphasizing the need for harmonized measurements, reference intervals, and reference change values for future clinical applications [2].

This molecular resolution is particularly relevant in dietary intervention studies, where standard lipid measures may remain unchanged even when lipidomics reveals coordinated shifts in specific lipid species [3].

Despite this potential, the biological interpretation of plasma lipidomics remains rarely straightforward. Current workflows often produce lists of altered lipid species, fold changes, p-values, multivariate loadings, or enrichment results. Although statistically informative, these outputs are not always easy to translate into nutritional or metabolic meaning. Changes in triacylglycerols, phosphatidylcholines, ceramides, plasmalogens, ether lipids, or polyunsaturated fatty acid-containing lipids may reflect distinct aspects of lipid storage, membrane remodelling, inflammatory tone, oxidative protection, or cardiometabolic stress [4, 5]. However, these meanings are usually distributed across many individual lipid species and are difficult to summarize in a form that is immediately useful for nutrition or clinical researchers. Beyond interpretation, an additional challenge concerns the comparability of lipidomic results across studies. In well-designed case-control cohorts, demographic and pre-analytical sources of variability can be mitigated through appropriate matching, balancing, and statistical adjustment. Nevertheless, findings based on individual lipid species may remain difficult to compare across independent datasets and analytical platforms. This limitation supports the use of biologically informed lipid indices that summarize distributed molecular changes into more robust nutritional and metabolic patterns.

These challenges are particularly relevant in human nutrition studies, where dietary interventions may induce subtle but coordinated changes in lipid classes, fatty acyl composition, saturation, chain length, ether lipid abundance, and neutral-to-polar lipid balance. Similarly, metabolic phenotypes such as insulin resistance involve broad lipidomic remodelling rather than isolated changes in single lipid markers [6]. There is therefore a need for computational tools that convert lipidomic matrices into biologically meaningful lipid signatures aligned with nutritional and metabolic interpretation.

To address this gap, we developed the Nutritional-Metabolic Lipid Profile (NMLP) module within LipidOne [7]. NMLP is designed as an interpretative layer that translates plasma lipidomics data into lipid indices, higher-order nutritional-metabolic categories, visual summaries, and concise interpretative statements.The rationale of NMLP is that biologically related lipid changes can be more informative when summarized as functional descriptors rather than interpreted only as isolated lipid species. For example, a coordinated increase in triacylglycerol-related indices and neutral-to-polar lipid ratios may suggest a shift toward lipid storage and energy-load features, whereas reduced ether lipid or plasmalogen-related indices may indicate altered membrane composition or lower oxidative-protective lipid components. By organizing lipidomic variation into interpretable dimensions, NMLP aims to make lipidomics more accessible to researchers and clinicians interested in nutritional metabolism, cardiometabolic health, and precision nutrition. In this study, we describe the NMLP framework and evaluate its performance in three human plasma lipidomics datasets relevant to nutrition and metabolic health: a randomized crossover feeding study comparing low- and high-glycemic-load dietary patterns [3], a eucaloric high-fat diet intervention in normal-weight women [8], and a large public lipidomics dataset stratified by insulin sensitivity [6]. These case studies were selected to test whether NMLP can support interpretation across controlled dietary interventions and clinically relevant metabolic phenotypes.

## 2. Methods

### 2.1. NMLP framework

NMLP was implemented within LipidOne as a rule-based workflow for deriving nutritional-metabolic descriptors from plasma lipidomics matrices. The workflow links lipid identifiers and abundance values to predefined lipid pools and indices reflecting class distribution, neutral-to-polar lipid balance, structural lipid organization, fatty acid remodelling, saturation, chain length distribution, ether lipid representation, and omega-3/essential fatty acid status. The input consists of a lipidomics matrix in which samples are arranged in columns and lipid variables in rows, together with a label row defining the experimental or clinical groups. Lipid identifiers are parsed to extract lipid class information and, when available, fatty acyl composition, according to established lipid classification and shorthand annotation principles [9]. NMLP then reconstructs quantitative lipid pools by summing species belonging to selected classes, subclasses, or fatty acyl features, and uses these pools to calculate nutritional-metabolic indices.

The resulting index matrix represents each sample by a reduced set of biologically interpretable variables. These variables are not intended to replace conventional lipid species-level analysis, but to complement it by providing a higher-level view of lipidomic remodelling.

### 2.2. Lipid harmonization, index calculation and category assignment

Because lipidomics datasets differ in nomenclature and annotation depth, LipidOne includes a lipid harmonization step before index calculation. Lipid identifiers are standardized to recognize lipid class, total carbon number, total number of double bonds, and, when available, individual fatty acyl chains. The NMLP module distinguishes between molecular-species-level annotations, such as PC 16:0/18:2, and sum-composition-level annotations, such as PC 34:2, in line with standardized lipid shorthand notation and reporting recommendations for MS-derived lipid structures [9, 10].

Depending on annotation depth, NMLP computes either the full set of indices or only those compatible with the available structural information. When chain-level data are available, the module calculates indices related to saturation, unsaturation, chain length, omega-3 and omega-6 representation, essential fatty acid status, and fatty acid remodelling. When only sum-composition data are available, the analysis is restricted to class-level or composition-compatible indices, such as neutral-to-polar lipid balance, structural lipid representation, and selected cardiometabolic lipid ratios. This avoids overinterpretation of datasets with limited structural resolution. The use of index-based lipidomic descriptors extends a strategy previously implemented in LipidOne for Functional Lipid Analysis, where lipid-derived indices were used to summarize complex lipidomic changes into biologically interpretable functional readouts [11].

Calculated indices are assigned to five higher-order nutritional-metabolic categories: Cardiometabolic Lipid Status, Fatty Acid Remodelling Status, Overall Lipid Quality, Oxidative Protection, and Omega-3 / Essential Fatty Acid Status. These categories were defined to organize lipidomic variation into domains relevant to nutrition and metabolic health. They summarize features related to neutral lipid accumulation, fatty acyl remodelling, lipid quality, ether/plasmalogen-associated oxidative protection, and essential fatty acid representation. The overall structure of the NMLP framework, from lipidomics input data to nutritional-metabolic interpretation, is summarized in Fig. 1. The full list of NMLP indices, formulas and category assignments is provided in Supplementary Table S1.

**Figure 1.**
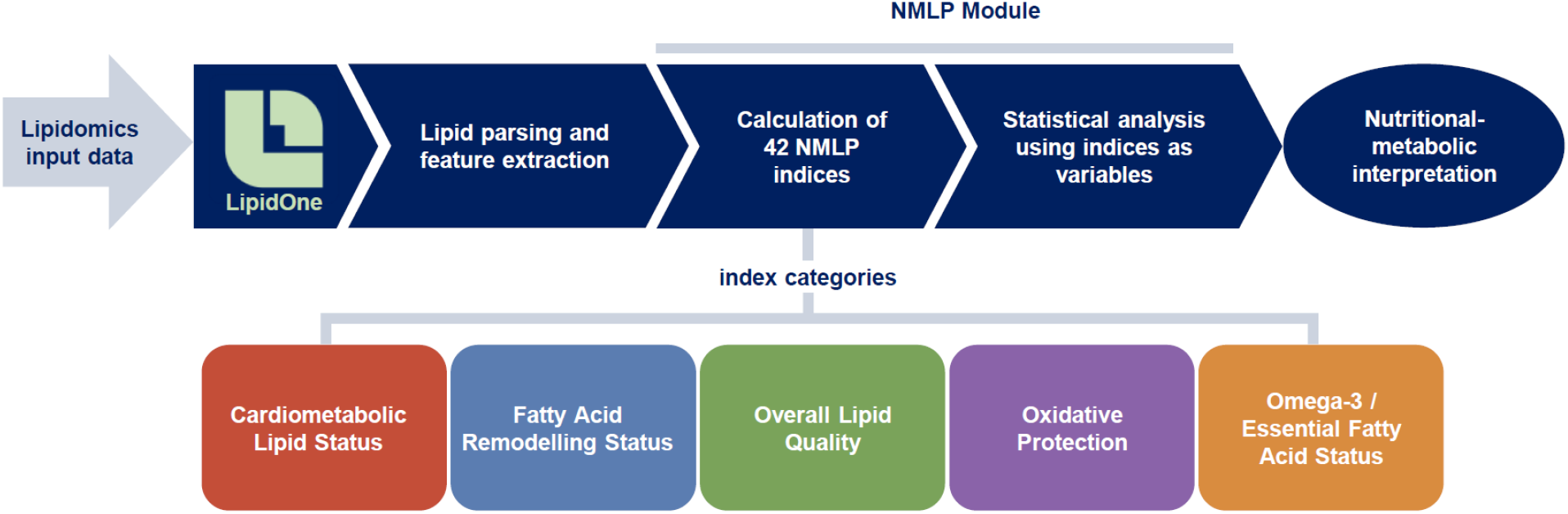
Overview of the Nutritional-Metabolic Lipid Profile (NMLP) module within LipidOne. Lipidomics input data are processed in LipidOne through lipid parsing and feature extraction, followed by calculation of NMLP indices and statistical analysis using indices as variables. The resulting outputs support nutritional-metabolic interpretation at both index and category levels. NMLP organizes lipidomic variation into five higher-order categories: Cardiometabolic Lipid Status, Fatty Acid Remodelling Status, Overall Lipid Quality, Oxidative Protection, and Omega-3 / Essential Fatty Acid Status.

### 2.3. Human lipidomics datasets

NMLP was evaluated and tested using three independent human plasma lipidomics datasets selected to represent complementary contexts of nutritional and metabolic research.

The first dataset was derived from a randomized controlled crossover feeding study comparing low- and high-glycemic-load dietary patterns, originally reported by Dibay Moghadam et al. and available through Metabolomics Workbench as ST001490 [3]. This dataset was selected because dietary glycemic load modified plasma lipidomic profiles despite limited changes in standard clinical lipid panel measures.

The second dataset was obtained from a one-month eucaloric high-fat diet intervention in normal-weight women, reported by Santoro et al. and available through Metabolomics Workbench as ST003044 [8]. It was used to evaluate whether NMLP could capture diet-induced lipid remodelling under controlled energy intake.

The third dataset was the longitudinal human plasma lipidomics study reported by Hornburg et al., available through Metabolomics Workbench as ST002081 [6]. This dataset was selected to test whether NMLP could identify functional lipidomic features associated with insulin resistance and reveal category-level heterogeneity within this metabolic phenotype.

For each dataset, lipidomic matrices were formatted according to the LipidOne input structure and processed through the NMLP workflow. Dataset characteristics are summarized in Table 1.

**Table 1.**
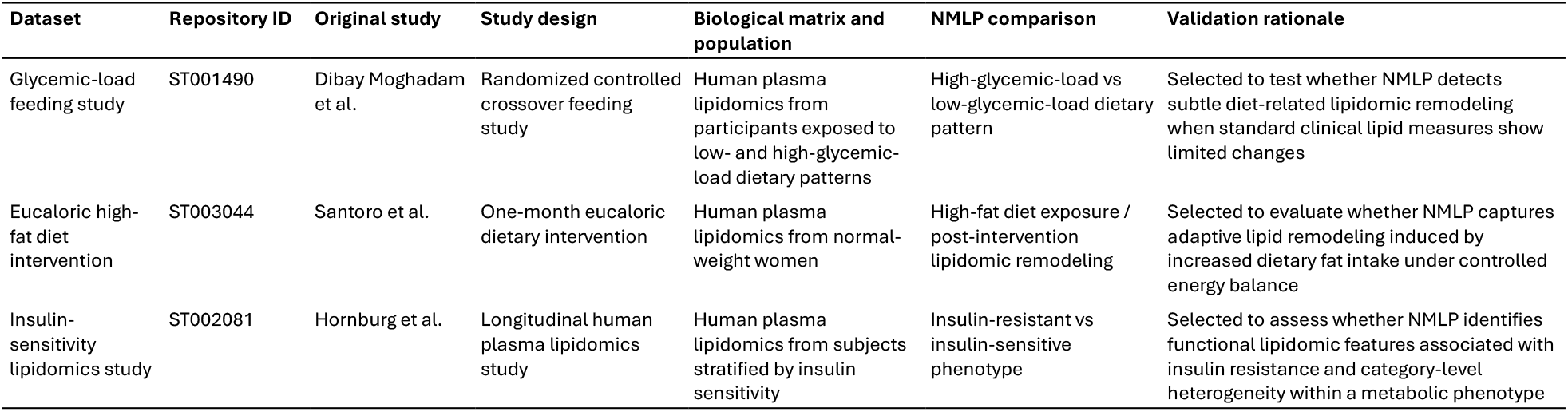
Characteristics of the public human plasma lipidomics datasets used to evaluate the NMLP module.

### 2.4. Statistical analysis and visualization

Statistical analyses were performed on the calculated NMLP index matrix. For two-group comparisons, index values were compared between groups, and the magnitude of change was summarized as log2 fold change. For each index, NMLP reports the direction of change, statistical significance, and effect magnitude. These values were used to generate volcano plots, bar plots, and interpretative tables. The rule-based interpretation table used to generate direction-dependent biological statements is provided in Supplementary Table S1.Category-level summaries were obtained by aggregating related indices within each nutritional-metabolic domain. These summaries were visualized using bar plots, radar plots, heatmaps, and sample-level profiles. Principal component analysis and clustering were performed on the index matrix to evaluate whether samples separated according to dietary intervention, metabolic phenotype, or functional lipidomic state. Hierarchical clustering was used to identify subgroups of samples with similar nutritional-metabolic profiles, particularly within insulin-resistant subjects.

Selected index changes were associated with concise biological statements according to their direction of change. In addition to group-level plots, NMLP generates three report-oriented outputs that summarize sample level deviations from a reference condition, dominant nutritional-metabolic categories, and the main indices contributing to the observed lipidomic phenotype as shown in Supplementary Report (Bar_Report, Lollipop_Report, Volcano_Report). These reports are intended to support clinical-nutritional interpretation and metabolic phenotyping, but not to provide automated diagnostic conclusions. All statistical and graphical outputs were generated within the LipidOne/NMLP workflow.

## 3. Results

### 3.1. NMLP generated interpretable nutritional-metabolic profiles across datasets

Across the three human plasma lipidomics datasets, NMLP transformed lipid species matrices into reduced index-based profiles that could be interpreted at both index and category levels. Instead of returning only individual lipid species, the module generated descriptors related to cardiometabolic lipid status, fatty acid remodelling, overall lipid quality, oxidative protection, and omega-3/essential fatty acid status. This structure allowed the same analytical framework to be applied to controlled dietary interventions and to a metabolic phenotype stratified by insulin sensitivity.

The index-based representation reduced the complexity of lipidomic interpretation while preserving biologically relevant information. In all datasets, NMLP outputs provided a functional summary of lipidomic remodelling, highlighting whether the main changes were related to neutral lipid storage, structural lipid organization, fatty acid composition, ether/plasmalogen features, or essential fatty acid-related profiles. The graphical outputs further supported this interpretation by combining index-level changes with category-level summaries and sample-level clustering.

### 3.2. Glycemic-load intervention: metabolic lipid shifts beyond standard lipid panels

In the randomized crossover glycemic-load feeding study, NMLP identified metabolic lipidomic changes between dietary conditions that were not immediately evident from standard clinical lipid measures. The index-level analysis showed that the high-versus low-glycemic-load contrast involved a limited but interpretable set of changes, including cardiometabolic lipid status, oxidative protection, and fatty acid remodelling indices (Fig. 2). Category-level summarization indicated that the response was mainly driven by oxidative-protection and cardiometabolic lipid-status features, whereas omega-3/essential fatty acid status contributed minimally to the contrast (Fig. 3). This was relevant because the original study reported limited changes in conventional lipid panel variables despite detectable lipidomic differences. NMLP therefore helped translate species-level dietary effects into broader functional descriptors, supporting the use of plasma lipidomics to capture subtle metabolic responses to dietary carbohydrate quality.

**Figure 2.**
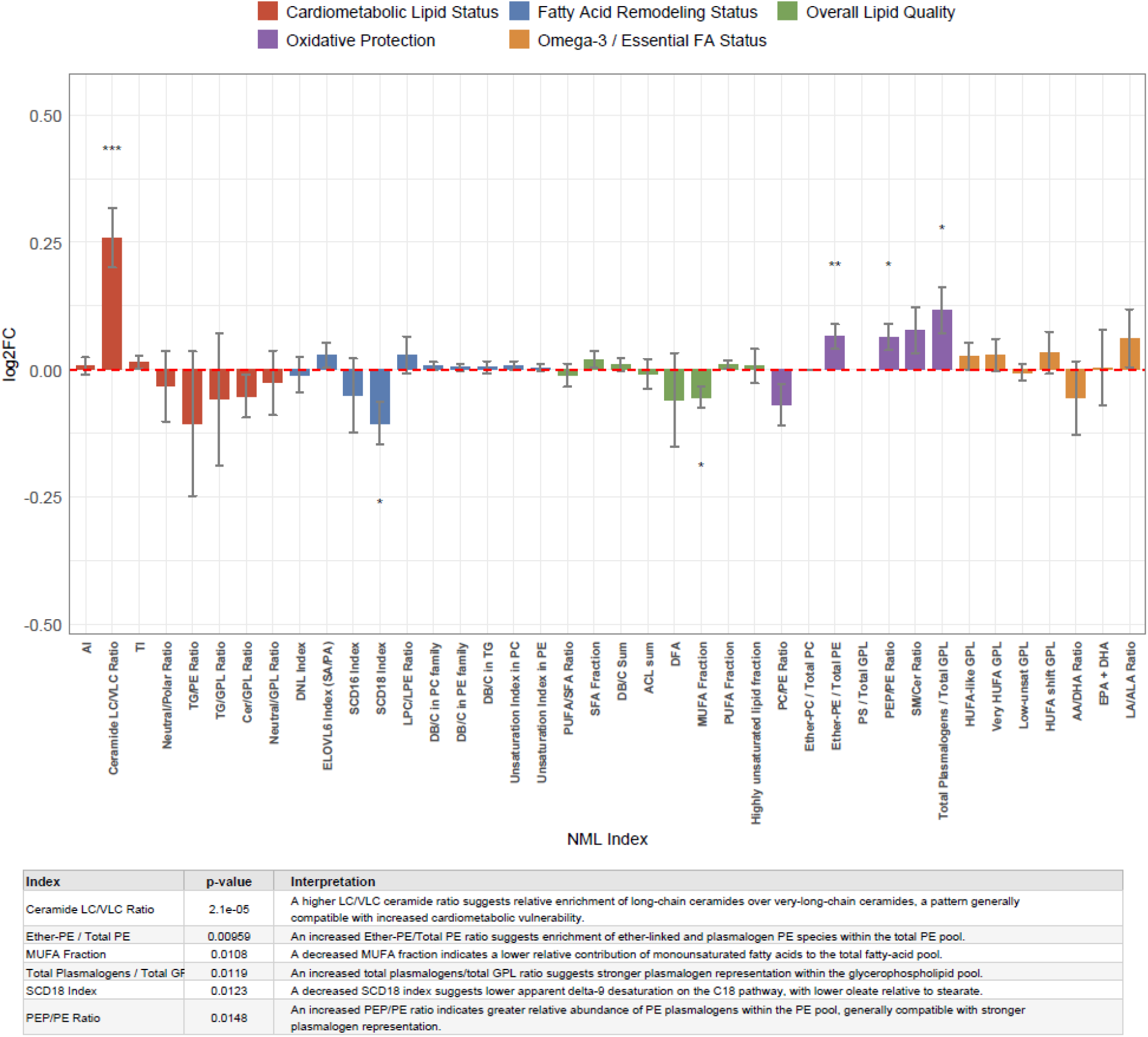
Index-level NMLP profile of the glycemic-load feeding study. NMLP index-level comparison between high-glycemic-load and low-glycemic-load dietary conditions in the ST001490 dataset (n = 80 participants/samples per condition). Bars show log2 fold-change values for individual NMLP indices and are coloured according to the five nutritional-metabolic categories. Error bars indicate the standard error of the mean (SEM). Asterisks indicate nominal statistical significance (p < 0.05, **p < 0.01, **p < 0.001). The table reports significant indices together with the corresponding p-values and direction-dependent biological interpretations generated from the NMLP interpretative rule set.

**Figure 3.**
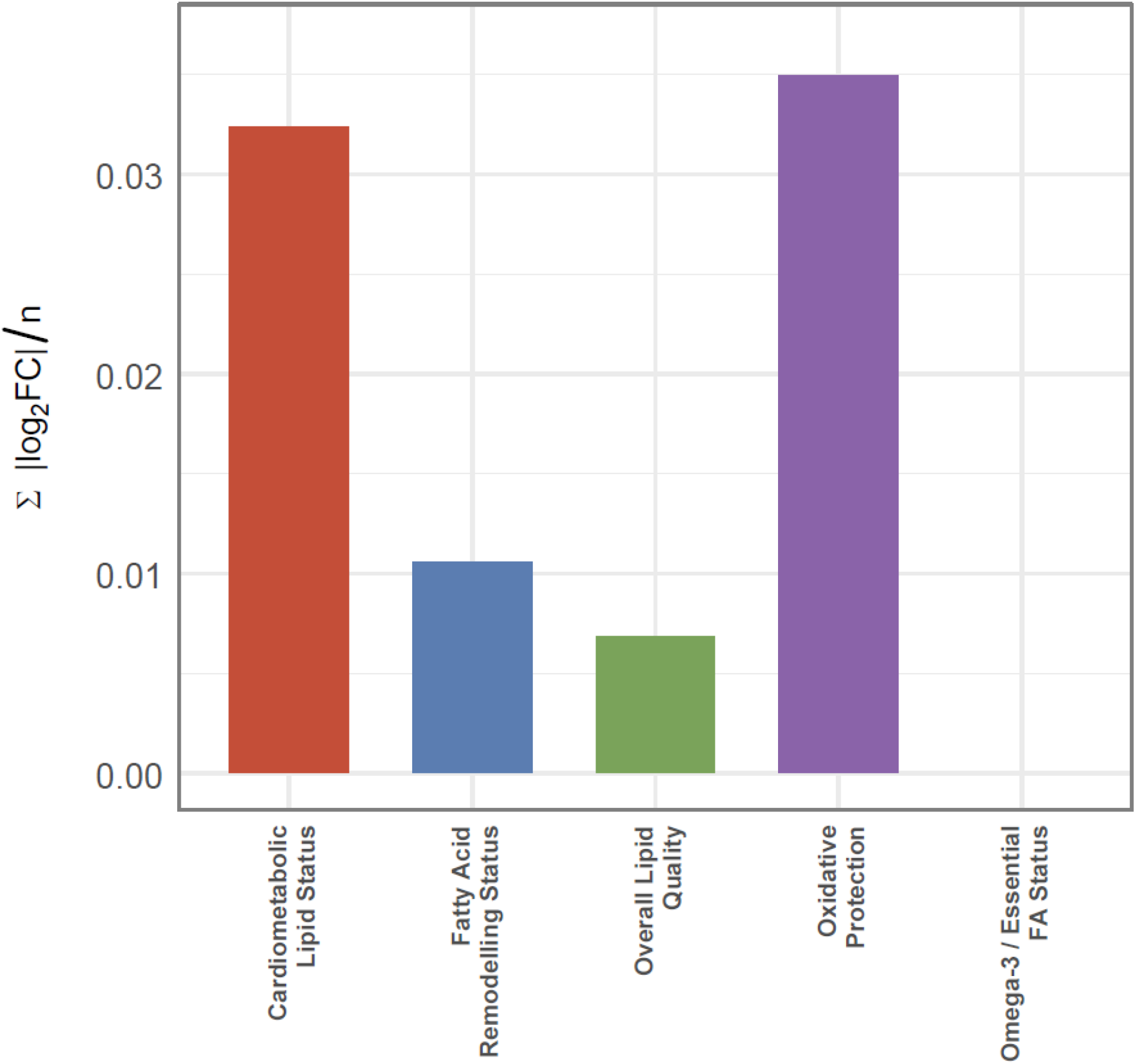
Category-level summary of the glycemic-load NMLP response. Normalized category-level summary of the high-glycemic-load versus low-glycemic-load contrast in the ST001490 dataset. Bars represent the aggregate contribution of significant NMLP index changes within each nutritional-metabolic category, normalized by the number of available indices in that category. The plot summarizes the relative involvement of Cardiometabolic Lipid Status, Fatty Acid Remodelling Status, Overall Lipid Quality, Oxidative Protection, and Omega-3 / Essential Fatty Acid Status in the dietary lipidomic response.

### 3.3. High-fat diet intervention: trajectory of metabolic lipid remodelling

In the eucaloric high-fat diet intervention, NMLP captured temporal lipid remodelling across pre-diet, on-diet and post-diet states. Category-level trajectories showed a transient increase in cardiometabolic lipid-status indices during the diet, followed by partial attenuation after the intervention, while oxidative-protection-related indices increased more clearly in the post-diet phase (Fig. 4A). Index-level trajectories within the Oxidative Protection category revealed heterogeneous responses among ether-, plasmalogen- and phospholipid-related indices (Fig. 4B). These results suggest that NMLP can describe diet-induced lipid remodelling as a dynamic process rather than as a single static contrast. Additional category-specific trajectories are shown in Supplementary Fig. 4. Reports related to the study is shown in Supplementary Report.

**Figure 4.**
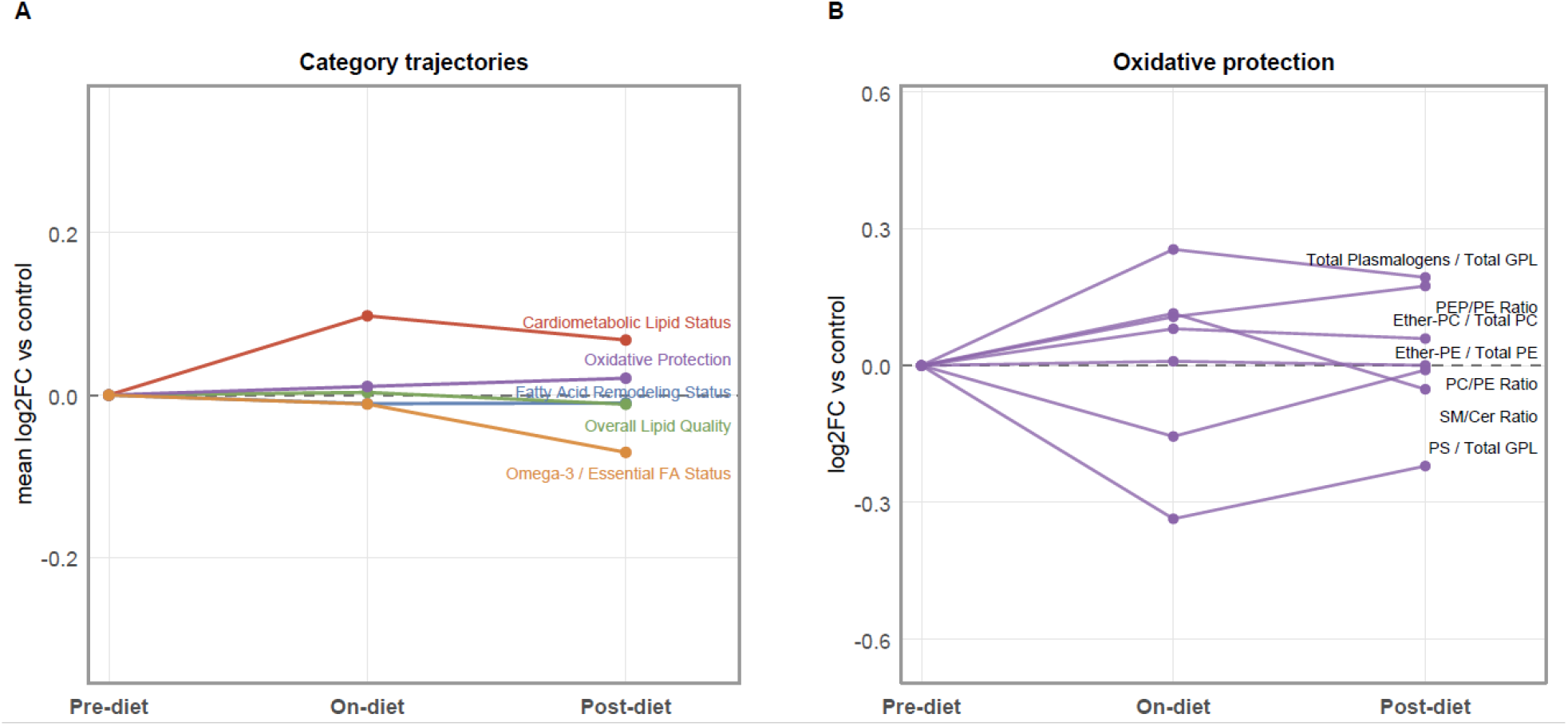
Temporal NMLP trajectories during and after a eucaloric high-fat diet intervention. (A) Mean category-level NMLP trajectories across pre-diet, on-diet, and post-diet conditions in the ST003044 dataset, expressed as log2 fold change relative to the pre-diet state. Lines represent the five NMLP nutritional-metabolic categories. (B) Representative index-level trajectories within the Oxidative Protection category, showing heterogeneous temporal responses of ether-, plasmalogen-, and phospholipid-related indices during and after the intervention.

### 3.4. Insulin-sensitivity dataset: neutral lipid storage, reduced structural features and category-level heterogeneity

In the insulin-sensitivity dataset, NMLP revealed a pronounced storage-oriented lipid phenotype in insulin-resistant subjects. The strongest changes involved neutral-to-polar lipid balance, TG/GPL ratio, TG/PE ratio and related cardiometabolic lipid-status indices, indicating increased neutral lipid storage relative to structural phospholipid pools (Fig. 5). This pattern was accompanied by changes in lipid quality and oxidative-protection-related indices, suggesting that insulin resistance was associated not only with increased storage lipids, but also with broader remodelling of structural and ether-linked lipid pools. Category-level clustering further separated insulin-resistant subjects into two subgroups with distinct nutritional-metabolic profiles (Fig. 6A). Cluster-level summaries showed divergent patterns across cardiometabolic lipid status, fatty acid remodelling, overall lipid quality, oxidative protection and omega-3/essential fatty acid status (Fig. 6B). Extended clustering outputs, including heatmap and dendrogram visualizations, are provided in Supplementary Fig. 6. These results indicate that individuals classified within the same insulin-resistant phenotype may carry different functional lipidomic states, supporting the potential use of NMLP for more refined metabolic phenotyping.

**Figure 5.**
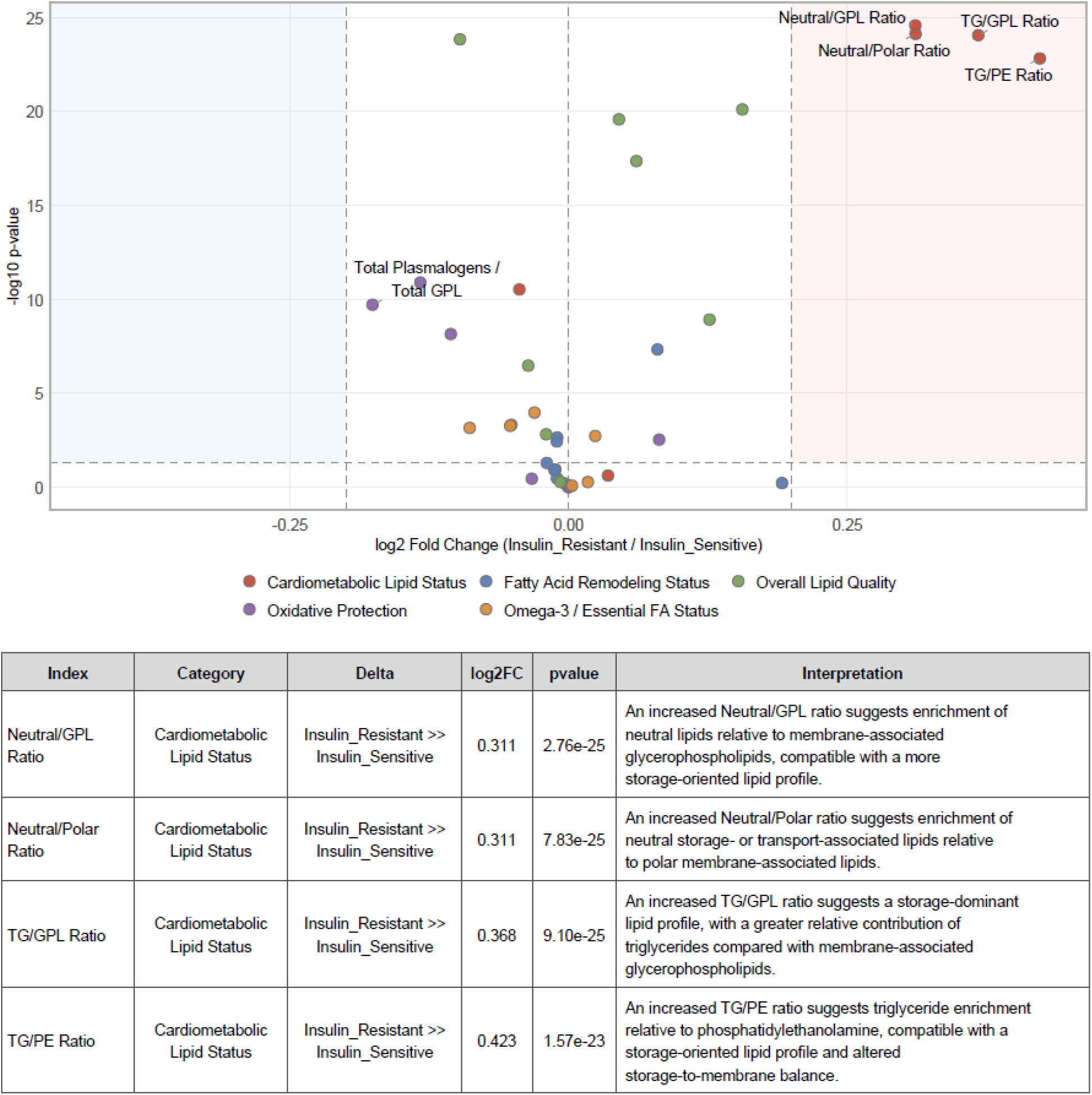
NMLP identifies a storage-oriented lipid phenotype in insulin-resistant subjects. Volcano plot of NMLP indices comparing insulin-resistant and insulin-sensitive subjects in the ST002081 dataset. Points represent individual NMLP indices, coloured according to nutritional-metabolic category. The x-axis shows log2 fold change and the y-axis shows −log10 p-value. Dashed lines indicate the fold-change and significance thresholds. Labelled points highlight the most relevant altered indices. The interpretation table summarizes significant storage- and cardiometabolic-related indices, their direction of change, p-values, and corresponding NMLP biological interpretations.

**Figure 6.**
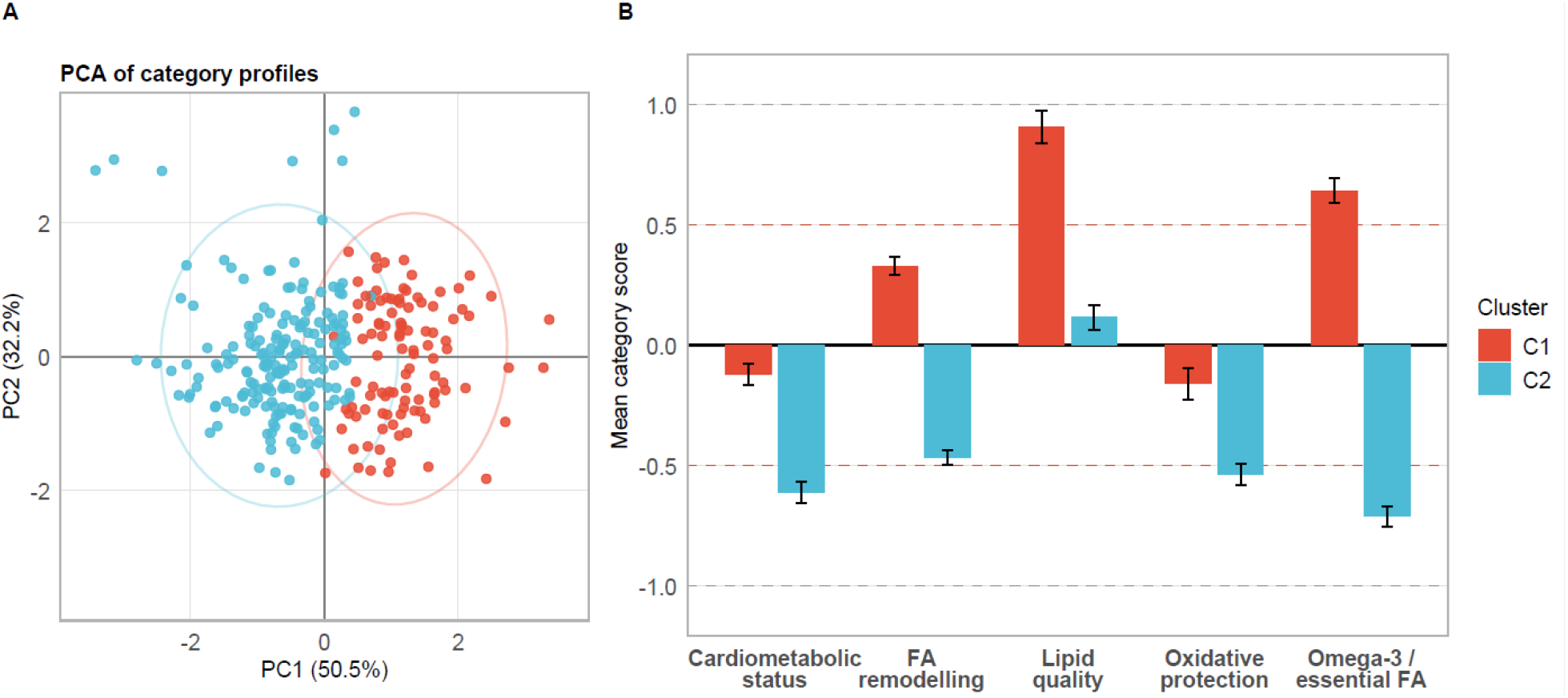
Category-level heterogeneity within insulin-resistant subjects. (A) Principal component analysis of NMLP category profiles within insulin-resistant subjects from the ST002081 dataset. Colours indicate hierarchical clustering groups, and ellipses represent cluster envelopes. (B) Mean NMLP category profiles of the two insulin-resistant clusters. Bars show cluster mean category scores and error bars indicate SEM. The two clusters display divergent nutritional-metabolic profiles across cardiometabolic lipid status, fatty acid remodelling, lipid quality, oxidative protection, and omega-3/essential fatty acid status.

## 4. Discussion

In this study, we developed and evaluated the Nutritional-Metabolic Lipid Profile module of LipidOne as an interpretative framework for human plasma lipidomics in nutrition and metabolic health research. Across three independent datasets, NMLP converted complex lipid species matrices into index-based and category-level profiles that captured biologically plausible dietary and metabolic lipidomic changes. The main contribution of the module is not to replace species-level lipidomics, but to add an intermediate interpretative layer between molecular lipid measurements and nutritional-metabolic meaning.The three validation datasets illustrate complementary uses of the framework. In the glycemic-load feeding study, NMLP summarized lipidomic changes that were not captured by standard clinical lipid measures, supporting the idea that plasma lipidomics can reveal subtle diet-related metabolic responses beyond conventional lipid panels. In the eucaloric high-fat diet intervention, the module described temporal lipid remodelling compatible with metabolic adaptation to increased dietary fat exposure, without necessarily implying pathological dysregulation. In the insulin-sensitivity dataset, NMLP revealed a stronger metabolic phenotype characterized by increased neutral lipid storage and energy-load features, together with changes in structural and ether-lipid-related indices. This pattern is consistent with the broader concept that insulin resistance involves coordinated lipidomic remodelling rather than isolated changes in single lipid markers.An important feature of the NMLP approach is that it preserves biological directionality while reducing dimensionality. In dietary studies, individual lipid species may change modestly and inconsistently, whereas coordinated shifts across lipid classes, chain features or lipid ratios may reveal a more coherent metabolic response. By summarizing these distributed signals into predefined indices and categories, NMLP may help distinguish adaptive dietary remodelling from potentially adverse metabolic lipid patterns. This is particularly relevant for clinical nutrition, where interventions are often expected to induce subtle, system-wide changes rather than large alterations in single lipid markers.

A key advantage of NMLP is that it reorganizes lipidomic information into domains relevant to nutrition and metabolic health. By grouping indices into cardiometabolic lipid status, fatty acid remodelling, overall lipid quality, oxidative protection, and omega-3/essential fatty acid status, the module provides a compact vocabulary for describing plasma lipidomic phenotypes. This may be useful in clinical nutrition studies, where the biological signal is often distributed across many lipid species and may be difficult to interpret from lists of significant molecules alone. The report-oriented outputs may further support the translation of lipidomic results into structured clinical-nutritional summaries, for example by highlighting the dominant altered categories and the indices most responsible for sample- or group-level deviations from a reference profile.

The index-based structure also allows multivariate and clustering analyses to be interpreted in terms of nutritional-metabolic dimensions rather than only as abstract statistical separation.

The clustering results in the insulin-resistance dataset further suggest that NMLP may support more refined metabolic phenotyping. Insulin-resistant subjects did not appear as a single uniform lipidomic group but showed heterogeneous category-level profiles. This observation is relevant from a precision-nutrition perspective, because individuals sharing the same clinical classification may differ in whether their lipidomic profile is dominated by storage-oriented lipids, reduced structural lipid quality, altered oxidative-protection features, or essential fatty acid imbalance. Although these findings require validation in prospective and intervention studies, they indicate that NMLP-derived profiles may help stratify metabolic states beyond conventional group labels.

It is important to emphasize that the strength of NMLP does not rely on the diagnostic value of any single index. Some indices are directly derived from established lipidomic or cardiometabolic literature, whereas others adapt nutritional fatty-acid concepts or class-level lipid balances to LC–MS lipidomics data. Their value lies primarily in their combined interpretation: when multiple independent descriptors move coherently across biological conditions, they provide a structured and biologically plausible readout of nutritional-metabolic lipid remodelling.

Several limitations should be considered. First, NMLP depends on the quality and structural resolution of the input lipidomics data. Some indices require molecular-species or chain-level annotation and cannot be fully computed from sum-composition data. Second, the biological interpretation of lipid indices remains context-dependent: a change in a lipid ratio may have different implications depending on the population, diet, disease state, analytical platform, and lipid classes detected. Third, the present study reanalysed existing datasets and was designed to evaluate interpretative performance rather than to establish clinical thresholds or diagnostic cut-offs. Finally, the biological statements generated by NMLP should be considered as structured interpretative annotations designed to orient data analysis, not as automated clinical conclusions.

Despite these limitations, NMLP provides a practical approach for translating plasma lipidomics into biologically meaningful descriptors. By combining lipid harmonization, index calculation, category-level organization, visualization, and direction-based interpretation, the module may help nutrition and clinical researchers extract metabolic information from complex lipidomic datasets. Future work should validate NMLP-derived profiles in larger prospective cohorts, controlled dietary interventions, and studies with clinical outcomes, and should evaluate whether these profiles improve metabolic risk stratification or personalized dietary recommendations.

In conclusion, NMLP extends LipidOne by providing an index-based framework for the nutritional-metabolic interpretation of human plasma lipidomics. By transforming lipid species data into biologically oriented indices, category-level summaries, visual outputs, and concise interpretative statements, the module facilitates the visualization, comparison, and interpretation of lipidomic profiles across dietary and metabolic contexts. This approach may support the integration of lipidomics into clinical nutrition, metabolic phenotyping, and precision nutrition research.

## Acknowledgements

We would like to thank Matteo Boschi (UMMON.it) for his invaluable cooperation in developing the LipidOne 2.5 website front-end and coding its back-end interactions with the R scripts.

## Author contributions

Conceptualisation: RMP, DFM and HBRA. Software development: RMP. Data analysis: DFM. Data validation: DFM and HBRA. Data interpretation: RMP, DFM and HBRA. Writing—original draft: DFM and RMP. Writing— review and editing: RMP, DFM and HBRA. All authors read and approved the final manuscript.

## Data availability

The original lipidomics datasets analysed in this study are publicly available through Metabolomics Workbench under the repository identifiers ST001490, ST003044 and ST002081. For reproducibility, the processed and harmonized input matrices used for the NMLP analyses are provided as Supplementary Material. These matrices include lipid shorthand harmonization, merging of duplicated lipid entries, group-label formatting, and dataset-specific group selection applied before analysis. The NMLP module is freely accessible as part of the LipidOne platform.

