## Supplementary Table S1 for "Nutritional-Metabolic Lipid Profiling with LipidOne for plasma lipidomics interpretation in metabolic health"

### Nutritional-Metabolic Lipid Profile indices, formulas, interpretive rules, biological rationale, and supporting references.

This table reports the complete set of indices implemented in the NMLP module, including their category assignment, index code, display name, formula, direction-specific interpretive phrases, biological explanation, and supporting references.

| Category | Index Code | Index Name | Formula | Interpretive phrase if increased | Interpretive phrase if decreased | Biological explanation | Ref. |
|---|---|---|---|---|---|---|---|
| Cardiometabolic Lipid Status | AI | Atherogenic Index (AI) | [C12:0 + (4×C14:0) + C16:0] / [ΣMUFA + Σn-6 PUFA + Σn-3 PUFA] | Higher AI indicates a more atherogenic fatty-acid profile, with greater weight of C12:0, C14:0 and C16:0 relative to unsaturated fatty acids. | Lower AI indicates a less atherogenic fatty-acid profile, with a greater relative contribution of unsaturated fatty acids. | The Atherogenic Index (AI) expresses the relationship between the principal saturated fatty acids considered pro-atherogenic, particularly lauric (C12:0), myristic (C14:0), and palmitic (C16:0) acids, and the unsaturated fatty acid fraction, which is regarded as relatively protective with respect to lipid deposition and plaque formation. A higher AI indicates a fatty acid pattern in which the relative contribution of these saturated fatty acids predominates over that of unsaturated fatty acids, consistent with a more atherogenic and less cardioprotective lipid profile, whereas a lower AI reflects a more favourable balance of unsaturated to pro-atherogenic saturated fatty acids and is therefore interpreted as more desirable for cardiovascular health. AI is widely used in food lipid studies to compare the cardiovascular quality of fats across animal-derived foods and processed lipid matrices. | 1, 2 |
| | Ceramide_LC_VLC_Ratio | Ceramide LC/VLC Ratio | (CER 16:0 + CER 18:0) / (CER 22:0 + CER 24:0) | A higher LC/VLC ceramide ratio suggests relative enrichment of long-chain ceramides over very-long-chain ceramides, a pattern generally compatible with increased cardiometabolic vulnerability. | A lower LC/VLC ceramide ratio suggests relative enrichment of very-long-chain ceramides over long-chain ceramides, a pattern generally compatible with a more favourable ceramide chain-length balance. | The Ceramide LC/VLC Ratio (Cer16:0+Cer18:0)/(Cer22:0+Cer24:0) is interpreted in NMLP as a composite ceramide chain-length balance descriptor. It summarizes the relative predominance of long-chain ceramides over very-long-chain ceramides, following the biological rationale that shorter long-chain ceramide species, particularly Cer 16:0 and Cer 18:0, have often been associated with lipotoxicity, impaired metabolic homeostasis, and adverse cardiometabolic outcomes, whereas higher very-long-chain/long-chain ceramide ratios have been linked to more favourable cardiovascular risk profiles. Community-based studies have reported inverse associations of C24:0/C16:0 and C22:0/C16:0 ceramide ratios with incident coronary heart disease, heart failure, and all-cause mortality, supporting the interpretation that a shift toward Cer 16:0 and Cer 18:0 relative to Cer 22:0 and Cer 24:0 reflects a less favourable ceramide remodelling pattern. In addition, dietary pattern studies indicate that circulating ceramide profiles and ceramide ratios are responsive to diet quality, including Mediterranean-style dietary patterns. Accordingly, in NMLP a higher LC/VLC ratio is interpreted as a ceramide signature more closely aligned with cardiometabolic risk, whereas a lower value indicates a relatively greater contribution of very-long-chain ceramides and a more favourable ceramide chain-length profile. | 3, 4, 5 |
| | TI | Thrombogenicity Index (TI) | TI = (C14:0 + C16:0 + C18:0) / [(0.5×ΣMUFA) + (0.5×Σn-6 PUFA) + (3×Σn-3 PUFA) + (n-3/n-6)] | Higher TI suggests a more thrombogenic fatty-acid pattern, with greater relative contribution of selected saturated fatty acids and lower protective unsaturated components. | Lower TI suggests a less thrombogenic fatty-acid pattern, with a more favourable contribution of MUFA and PUFA, especially n-3 PUFA. | The Index of Thrombogenicity (IT) is a composite fatty acid index specifically designed to estimate the theoretical thrombogenic potential of a lipid profile and is therefore best interpreted as a marker of cardiovascular risk orientation. The logic of the index: the numerator captures fatty acids considered relatively more pro-thrombogenic, mainly selected saturated fatty acids, whereas the denominator weighs fatty acids regarded as more anti-thrombogenic, especially MUFA, n-6 PUFA, and most strongly n-3 PUFA, thereby summarizing the balance between lipid components thought to influence platelet aggregation, eicosanoid balance, thrombus formation, and cardiovascular health. From a nutritional perspective, a lower TI is generally considered more favourable because it reflects a fat profile less dominated by thrombogenic saturated fatty acids and relatively richer in unsaturated fatty acids with protective cardiovascular associations; for this reason, TI is widely used to compare the nutritional quality of milk, dairy products, meats, fish, oils, and other foods. | 1, 2, 6 |
| | TG_GPL_Ratio | TG/GPL | Σ total triglycerides / Σ total glycerophospholipids | An increased TG/GPL ratio suggests a storage-dominant lipid profile, with a greater relative contribution of triglycerides compared with membrane-associated glycerophospholipids. | A decreased TG/GPL ratio suggests a lower relative contribution of triglycerides compared with glycerophospholipids, compatible with a less storage-dominant and more membrane-oriented lipid profile. | The TG/GPL ratio is a derived lipid-class balance descriptor that summarizes the relative contribution of neutral storage lipids and membrane-associated glycerophospholipids. A higher value indicates that the measured lipid pool is relatively enriched in triglycerides, consistent with a storage-oriented or lipid-accumulation phenotype, whereas a lower value indicates a relatively greater contribution of glycerophospholipids, compatible with a more membrane-oriented lipid organization. This index is not a standardized clinical or food-quality metric and should therefore be interpreted cautiously and in relation to the biological matrix analysed. However, its rationale is consistent with lipidomic studies showing that triglyceride-rich lipid signatures are associated with obesity-related and cardiometabolic alterations, while changes in phospholipid classes may reflect broader remodelling of lipid homeostasis. In NMLP, TG/GPL is therefore used as an exploratory descriptor of storage-dominant versus membrane-oriented lipid organization. | 7, 8 |
| | Cer_GPL_Ratio | Cer/GPL Ratio | ΣCer / total GPL | An increased Cer/GPL ratio suggests a higher relative ceramide load compared with membrane glycerophospholipids, compatible with a more lipotoxic and less favourable cardiometabolic lipid profile. | A decreased Cer/GPL ratio suggests a lower relative ceramide load compared with membrane glycerophospholipids, compatible with a less ceramide-enriched cardiometabolic lipid profile. | The Cer/GPL ratio, defined as total ceramides relative to total glycerophospholipids within the same lipidomic compartment, can be interpreted as a derived descriptor of relative ceramide load against the structural membrane phospholipid pool. This interpretation is biologically plausible because ceramides are bioactive sphingolipids increasingly linked to insulin resistance, inflammation, endothelial dysfunction, myocardial injury, lipotoxicity, and adverse cardiovascular outcomes, whereas glycerophospholipids constitute a major structural framework of cellular membranes and contribute to membrane adaptability and homeostasis. A higher Cer/GPL ratio is therefore consistent with a lipidome relatively more enriched in stress-related sphingolipid signaling compared with membrane-associated glycerophospholipid abundance, whereas a lower ratio suggests a less ceramide-enriched balance. This ratio should not be interpreted as a standardized clinical ceramide risk score, because established ceramide-based cardiometabolic risk markers often depend on specific ceramide species and species ratios. In NMLP, Cer/GPL is used as an exploratory cardiometabolic burden descriptor that summarizes the balance between ceramide-associated lipotoxic signaling and the glycerophospholipid membrane background. | 9, 10, 11 |
| | Neutral_GPL_Ratio | Neutral/GPL Ratio | (TG + DG + CE) / total GPL | An increased Neutral/GPL ratio suggests enrichment of neutral lipids relative to | A decreased Neutral/GPL ratio suggests a lower | The Neutral/GPL ratio is calculated as the sum of triglycerides, diacylglycerols, and cholesteryl esters divided by total glycerophospholipids within the same lipidomic compartment. This derived descriptor summarizes the relative contribution of neutral or storage-associated lipids compared with the glycerophospholipid membrane | 12, 13 |

| | | | | | | | |
|---|---|---|---|---|---|---|---|
| | | | | membrane-associated glycerophospholipids, compatible with a more storage-oriented lipid profile. | relative contribution of neutral lipids compared with glycerophospholipids, compatible with a less storage-dominant and more membrane-oriented lipid profile. | background. Glycerophospholipids are major structural components of biological membranes and contribute to membrane organization, adaptability, and homeostasis, whereas TG and CE mainly represent neutral lipid pools involved in energy storage and lipoprotein-core composition. DG is also included because it acts as both a metabolic intermediate and a bioactive lipid involved in lipid signaling and insulin-resistance-related pathways. Accordingly, a higher Neutral/GPL ratio may indicate a lipidome relatively enriched in neutral lipids compared with structural glycerophospholipids, compatible with a storage-oriented or lipid-accumulation phenotype. Conversely, a lower ratio suggests a relatively greater contribution of membrane-associated glycerophospholipids. This index is not a standardized clinical or nutritional metric and should be interpreted cautiously, especially because TG, DG, and CE represent biologically distinct neutral-lipid pools. In NMLP, Neutral/GPL is therefore used as an exploratory lipidomic balance descriptor of neutral-lipid enrichment versus glycerophospholipid membrane background. | |
| | Neutral_Polar_Ratio | Neutral/Polar Ratio | Neutral/Polar = Σ neutral lipids / Σ polar lipids | An increased Neutral/Polar ratio suggests enrichment of neutral storage- or transport-associated lipids relative to polar membrane-associated lipids. | A decreased Neutral/Polar ratio suggests a lower relative contribution of neutral lipids compared with polar membrane-associated lipids. | The Neutral/Polar lipid ratio summarizes the class-level distribution of a lipidome between neutral lipids, typically including triglycerides, diacylglycerols, cholesteryl esters and related storage- or transport-associated species, and polar lipids, mainly including glycerophospholipids and polar sphingolipids. Biochemically, this ratio is best interpreted as a broad descriptor of lipidomic architecture, because neutral lipids are major components of lipid droplets and lipoprotein cores, whereas polar lipids contribute to membrane structure, membrane remodelling, lipoprotein surfaces and lipid signaling. A higher Neutral/Polar ratio indicates a lipidome relatively enriched in neutral lipids compared with polar lipid classes, compatible with a more storage- or transport-oriented lipid profile. In human plasma or lipoprotein-related contexts, this pattern may be consistent with increased neutral-lipid accumulation, such as TAG-, DAG- or CE-rich profiles. Conversely, a lower ratio reflects a relatively greater contribution of polar membrane-associated lipids. This should not automatically be interpreted as metabolically favourable, because the biological meaning of the ratio depends on the sample type, lipid classes included, and whether the polar fraction is enriched in structural or bioactive stress-related lipids. In NMLP, Neutral/Polar is therefore used as an exploratory descriptor of neutral-lipid predominance versus polar lipid background. | 14, 15 |
| | TG_PE_Ratio | TG/PE Ratio | TG/PE = Σ triglycerides / Σ phosphatidylethanolamine | An increased TG/PE ratio suggests triglyceride enrichment relative to phosphatidylethanolamine, compatible with a storage-oriented lipid profile and altered storage-to-membrane balance. | A decreased TG/PE ratio suggests a lower relative triglyceride contribution compared with phosphatidylethanolamine, compatible with a less storage-dominant lipid profile. | The TG/PE ratio, defined as total triglycerides divided by total phosphatidylethanolamine within the same biological compartment, reflects the balance between neutral lipid storage and a major structural membrane phospholipid class. This ratio is therefore best interpreted as a derived storage-to-membrane lipid balance descriptor rather than as a classical fatty-acid quality index. Biologically, triglycerides represent a major neutral lipid class devoted to energy storage, particularly under conditions of nutrient excess, whereas phosphatidylethanolamine is one of the most abundant membrane phospholipids and contributes to membrane architecture, membrane curvature, protein stabilization, mitochondrial function, and cellular lipid homeostasis. A higher TG/PE ratio is consistent with a lipidome relatively shifted toward triglyceride accumulation compared with PE-associated membrane lipid abundance, a pattern that may be relevant in overnutrition, obesity-related metabolic disturbance, hepatic lipid overload, and steatosis. Conversely, a lower TG/PE ratio suggests a lower storage-lipid contribution relative to PE. This index is not a standardized clinical marker and should be interpreted together with the individual TG and PE levels, because an increased ratio may result from TG accumulation, PE depletion, or both. In NMLP, TG/PE is used as an exploratory cardiometabolic descriptor of neutral lipid storage relative to phosphatidylethanolamine-associated membrane background. | 16 |
| Fatty Acid Remodelling Status | DNL_Index | DNL Index | DNL index = 16:0 / 18:2n-6 | An increased DNL index suggests a stronger estimated de novo lipogenesis-related signature, with higher palmitate relative to linoleate. | A decreased DNL index suggests a weaker estimated de novo lipogenesis-related signature, with lower palmitate relative to linoleate. | The DNL index is intended to capture the relative contribution of endogenous fatty acid synthesis to the lipid pool by contrasting palmitate, the principal end product of de novo lipogenesis, with linoleic acid, an essential dietary fatty acid that cannot be synthesized de novo in humans. As such, it is best interpreted as an estimated de novo lipogenesis-related activity index rather than as a general fatty-acid quality metric. This index reflects a specific metabolic pathway through which excess carbohydrate-derived substrate can be converted into fatty acids, particularly in the liver. A higher DNL index is consistent with a greater estimated lipogenic contribution, whereas a lower DNL index suggests attenuation of the lipogenic signature. In nutritional-metabolic contexts, a lower value is often interpreted as metabolically favourable, especially when it accompanies reduced hepatic fat accumulation or improved lipid handling. However, the index should be interpreted cautiously because palmitate may also derive from dietary sources and linoleate reflects dietary intake and lipid-pool composition. In NMLP, the DNL index is therefore used as an exploratory fatty-acid remodelling descriptor of estimated de novo lipogenesis activity. | 17, 18, 19 |
| | ELOVL6_Index | ELOVL6 Index (SA/PA) | SA/PA = 18:0 / 16:0 | An increased ELOVL6 index suggests greater estimated elongation from palmitate to stearate. | A decreased ELOVL6 index suggests lower estimated elongation from palmitate to stearate. | The ELOVL6 index, commonly expressed as the stearic acid to palmitic acid ratio (18:0/16:0; SA/PA), is a product-to-precursor descriptor used to estimate the relative contribution of the C16:0 to C18:0 elongation step in fatty acid metabolism. Its rationale is linked to ELOVL6, an elongase involved in the conversion of palmitate (16:0) into stearate (18:0). A higher SA/PA ratio is consistent with greater estimated ELOVL6-related elongation activity, whereas a lower ratio is compatible with reduced apparent elongation and relative enrichment of palmitate compared with stearate. For this reason, the index is classified as a fatty acid remodelling marker rather than as a general lipid-quality index. In human studies, higher SA/PA ratios have been associated in specific contexts with more favourable metabolic features, including lower insulin-resistance-related markers and greater probability of diabetes remission after bariatric surgery. However, the ratio remains an indirect estimate and should be interpreted cautiously because 16:0 and 18:0 levels may also be influenced by diet, lipid class distribution, desaturation, sample type, and the biological compartment analysed. In NMLP, the ELOVL6 index is therefore used as an exploratory descriptor of estimated palmitate-to-stearate elongation activity. | 20, 21, 22 |
| | SCD16_Index | SCD16 Index | SCD16 = 16:1n-7 / 16:0 | An increased SCD16 index suggests higher apparent delta-9 desaturation on the C16 pathway, with greater palmitoleate relative to palmitate. | A decreased SCD16 index suggests lower apparent delta-9 desaturation on the C16 pathway, with lower palmitoleate relative to palmitate. | The SCD16 index is a product-to-precursor ratio used as a surrogate descriptor of apparent delta-9 desaturase activity on the C16 fatty acid pathway. It reflects the relative conversion of palmitic acid (16:0) into palmitoleic acid (16:1n-7), a reaction catalysed by stearoyl-CoA desaturase enzymes, particularly SCD1 in metabolic tissues. This ratio is informative because SCD1 introduces a cis double bond into saturated fatty acyl-CoAs, generating monounsaturated fatty acids that can be incorporated into triglycerides, phospholipids, cholesteryl esters, and other complex lipids. Accordingly, a higher SCD16 index is consistent with greater apparent C16 delta-9 desaturation and may reflect a more active endogenous desaturation/lipogenic phenotype, often associated with VLDL-triglyceride production, adiposity, hypertriglyceridemia, and metabolically less favourable lipid remodelling. However, the index remains an indirect estimate and should be interpreted cautiously because palmitoleate and palmitate levels may also be influenced by diet, lipid class distribution, lipoprotein | 23, 24, 25 |

| | | | | | | | |
|---|---|---|---|---|---|---|---|
| | | | | | | composition, sample type, and the correct assignment of the 16:1n-7 isomer. In NMLP, SCD16 is therefore used as an exploratory fatty-acid remodelling descriptor of apparent C16 delta-9 desaturation activity. | |
| | SCD18_Index | SCD18 Index | SCD18 = 18:1n-9 / 18:0 | An increased SCD18 index suggests higher apparent delta-9 desaturation on the C18 pathway, with greater oleate relative to stearate. | A decreased SCD18 index suggests lower apparent delta-9 desaturation on the C18 pathway, with lower oleate relative to stearate. | The SCD18 index, defined as 18:1n-9 / 18:0, is a product-to-precursor ratio used as a surrogate descriptor of apparent delta-9 desaturase activity on the C18 fatty acid pathway. It reflects the relative conversion of stearic acid (18:0) into oleic acid (18:1n-9), a reaction catalysed by stearoyl-CoA desaturase enzymes, particularly SCD1 in metabolic tissues. This ratio is informative because oleate is a major monounsaturated fatty acid incorporated into triglycerides, phospholipids, cholesteryl esters, and other complex lipids. A higher SCD18 index is therefore consistent with greater apparent C18 delta-9 desaturation and may reflect endogenous desaturation/lipogenic remodelling. However, compared with SCD16, this index is particularly sensitive to dietary oleic acid intake and lipid class distribution, and should not be interpreted as a direct measurement of SCD1 activity. Nutritionally, SCD18 is best considered an index of diet-related and metabolic fatty acid remodelling rather than a general food-quality marker. In NMLP, SCD18 is used as an exploratory descriptor of apparent C18 delta-9 desaturation activity, to be interpreted together with dietary context, sample type, and the individual levels of 18:1n-9 and 18:0. | 23, 24 |
| | DB_C_PC | DB/C_PC | total PC-family double bonds / total PC-family carbons | Higher DB/C in PC suggests a more unsaturated phosphatidylcholine pool relative to chain length. | Lower DB/C in PC suggests a less unsaturated phosphatidylcholine pool relative to chain length. | DB/C_PC, defined as total PC-family double bonds divided by total PC-family carbons, is a class-specific unsaturation-density index that quantifies how densely double bonds are distributed within the phosphatidylcholine pool. Phosphatidylcholine is one of the dominant structural glycerophospholipids in eukaryotic membranes and contributes to membrane organization, bilayer properties, and lipid–cholesterol interactions, while acyl-chain unsaturation is a key determinant of bilayer packing, order, and adaptability. By normalizing double-bond content to total PC carbons, these metric captures whether the PC fraction is becoming relatively more unsaturated or more saturated, with reduced confounding from simple chain-length shifts. Within this framework, a higher DB/C_PC indicates a PC pool enriched in unsaturated acyl architecture, compatible with a membrane composition that may favour lower packing, greater disorder, and increased fluidity, whereas a lower DB/C_PC indicates lower unsaturation density, generally compatible with tighter acyl-chain packing and greater membrane order. At the same time, greater phospholipid unsaturation also increases the availability of oxidizable double bonds. This index is therefore best regarded as a structural fatty-acid remodelling marker rather than as a uniformly favourable or unfavourable metabolic signal, because its biological meaning depends on the balance between membrane adaptability, lipid class context, cholesterol content, and oxidative environment. | 26 |
| | DB_C_PE | DB/C_PE | total PE-family double bonds / total PE-family carbons | Higher DB/C in PE suggests a more unsaturated phosphatidylethanolamin e pool relative to chain length. | Lower DB/C in PE suggests a less unsaturated phosphatidylethanola mine pool relative to chain length. | DB/C_PE, defined as total PE-family double bonds divided by total PE-family carbons, is a class-specific unsaturation-density index for the phosphatidylethanolamine pool. PE is a major membrane phospholipid with important roles in membrane curvature, fusion, organelle membrane remodelling, and mitochondrial membrane function. Because acyl-chain double bonds reduce lipid packing and influence bilayer order, this ratio summarizes how densely unsaturation is distributed within the PE fraction, independently of its absolute class abundance and with reduced confounding from simple chain-length shifts. A higher DB/C_PE indicates a PE pool enriched in unsaturated chains, compatible with a membrane composition that may favour flexibility and dynamic membrane remodelling. A lower DB/C_PE indicates lower unsaturation density, generally compatible with tighter packing and a less flexible PE architecture. The index may also reflect the relative incorporation of diet-related unsaturated fatty acids into the PE family. However, it should be interpreted as a fatty-acid remodelling descriptor rather than as a classical nutritional index or a direct measure of membrane fluidity. Its biological meaning is context-dependent: greater PE unsaturation may support membrane adaptability, but it can also increase susceptibility to oxidative damage, especially when polyunsaturated species are involved. | 27 |
| | DB_C_TG | DB/C_TG | total TG double bonds / total TG carbons | Higher DB/C in TG suggests a more unsaturated triglyceride pool relative to chain length. | Lower DB/C in TG suggests a less unsaturated triglyceride pool relative to chain length. | DB/C_TG, defined as total TG double bonds divided by total TG carbons, is a class-specific unsaturation-density index for the triglyceride pool. Triglycerides are primarily neutral lipids involved in energy storage, lipid droplet accumulation, and transport within triglyceride-rich lipoproteins, rather than structural membrane lipids. A higher DB/C_TG indicates a TG pool relatively richer in unsaturated acyl chains after normalization for total chain length, whereas a lower DB/C_TG indicates a TG pool with lower unsaturation density and relatively more saturated or less unsaturated acyl architecture. This index may reflect both dietary fat composition and endogenous fatty-acid remodelling, including desaturation and selective incorporation of fatty acids into TG species. In lipidomics studies, TG species with lower double-bond number have often been associated with less favourable metabolic phenotypes, whereas TG species with higher double-bond number have been associated with more favourable ones. However, the biological meaning of DB/C_TG remains context-dependent because unsaturation type matters: MUFA-, n-6 PUFA-, and n-3 PUFA-containing TGs may have different implications. Greater unsaturation may also increase the oxidation susceptibility of the TG fraction. In NMLP, DB/C_TG is therefore used as an exploratory descriptor of triglyceride unsaturation density rather than as a uniformly favourable or unfavourable metabolic index. | 28 |
| | UI_PC | UI_PC | total PC-family double bonds / total PC-family abundance | Higher UI in PC indicates a more unsaturated phosphatidylcholine pool. | Lower UI in PC indicates a less unsaturated phosphatidylcholine pool. | UI_PC is defined as total PC-family double bonds divided by total PC-family abundance and can be interpreted as a class-specific unsaturation index for the phosphatidylcholine pool. PC is one of the major structural phospholipids of eukaryotic membranes and contributes to membrane organization, bilayer packing, fluidity, and lipid–cholesterol interactions. A higher UI_PC indicates a PC pool enriched in more unsaturated species, compatible with a membrane composition that may favor lower packing order and greater fluidity. A lower UI_PC indicates a PC pool enriched in less unsaturated or more saturated species, compatible with tighter packing and a more ordered membrane environment. UI_PC may also reflect the relative incorporation of diet-related unsaturated fatty acids into the PC family. However, it should be interpreted as a fatty-acid remodelling descriptor rather than as a classical nutritional index or a direct measure of membrane fluidity. Compared with DB/C_PC, UI_PC is not normalized for total chain length and may therefore be more influenced by shifts in average PC chain length. Its interpretation remains context-dependent: greater PC unsaturation may support membrane adaptability, but it also increases the number of oxidizable double bonds. | 26 |
| | UI_PE | UI_PE | total PE-family double bonds / total PE-family abundance | Higher UI in PE indicates a more unsaturated phosphatidylethanolamin e pool. | Lower UI in PE indicates a less unsaturated phosphatidylethanola mine pool. | UI_PE is defined as total PE-family double bonds divided by total PE-family abundance and can be interpreted as a class-specific unsaturation index for the phosphatidylethanolamine pool. It reflects the average double-bond enrichment of PE species. PE is a major membrane phospholipid with important roles in membrane curvature, fusion, organelle membrane remodelling, and mitochondrial membrane function. A higher UI_PE indicates a PE pool enriched in more unsaturated species, compatible with a membrane composition that may favour greater flexibility and dynamic remodelling. A lower UI_PE indicates a PE pool enriched in less unsaturated or more | 27 |

| | | | | | | | |
|---|---|---|---|---|---|---|---|
| | | | | | | saturated species, compatible with tighter packing and lower membrane flexibility. UI_PE may also reflect the relative incorporation of diet-related unsaturated fatty acids into the PE family. However, it should be interpreted as a lipid remodelling descriptor rather than as a classical nutritional index or a direct measure of membrane fluidity. Compared with DB/C_PE, UI_PE is not normalized for total chain length and may therefore be more influenced by shifts in average PE chain length. Its biological meaning remains context-dependent: greater PE unsaturation may support membrane adaptability, but it may also increase susceptibility to oxidative damage, especially when polyunsaturated PE species are involved. | |
| | LPC_LPE_Ratio | LPC/LPE Ratio | LPC/LPE = ΣLPC / ΣLPE | An increased LPC/LPE ratio suggests a shift in lysophospholipid balance toward lysophosphatidylcholine relative to lysophosphatidylethanolamine. | A decreased LPC/LPE ratio suggests a shift in lysophospholipid balance toward lysophosphatidylethanolamine relative to lysophosphatidylcholine. | The LPC/LPE ratio reflects the remodelling balance between two major lysophospholipid pools generated from phosphatidylcholine and phosphatidylethanolamine through phospholipase-mediated deacylation and reacylation pathways involved in membrane phospholipid turnover, including the Lands cycle. Because LPC is commonly one of the most abundant lysophospholipid classes in plasma, whereas LPE is usually less abundant but biologically informative, a higher LPC/LPE ratio indicates a more LPC-dominant lysophospholipid profile, while a lower ratio indicates relative enrichment of the LPE-related compartment. This ratio should be interpreted primarily as a lysophospholipid remodelling descriptor rather than as a classical nutritional index. Some LPC species and LPC-rich profiles have been linked to vascular inflammation, oxidative stress, endothelial dysfunction, and atherosclerotic processes, but their biological meaning is species-, matrix-, and context-dependent. In NMLP, LPC/LPE is therefore used as an exploratory marker of lysophospholipid remodelling balance, describing whether a biological sample is relatively LPC-dominant or LPE-shifted, with potential pathophysiological relevance only when interpreted together with the specific molecular species and biological context examined. | 29, 59 |
| Omega-3 / Essential FA Status | AA_DHA_Ratio | AA/DHA Ratio | AA/DHA = 20:4n-6 / 22:6n-3 | An increased AA/DHA ratio suggests a shift toward arachidonic-acid predominance relative to DHA, compatible with a more AA-oriented inflammatory-signaling potential. | A decreased AA/DHA ratio suggests greater DHA representation relative to arachidonic acid, compatible with a more omega-3-oriented lipid balance. | The AA/DHA ratio, defined as 20:4n-6 divided by 22:6n-3, expresses the relative abundance of arachidonic acid and docosahexaenoic acid within a lipid matrix. It reflects the balance between two biologically important long-chain polyunsaturated fatty acids with partly distinct metabolic functions. AA is a major omega-6 constituent of membrane phospholipids and a precursor of numerous eicosanoids involved in cell signaling, vascular regulation, immune modulation, and inflammatory responses, whereas DHA is a major omega-3 structural component of neuronal and retinal membranes and contributes to membrane biophysical properties, neurodevelopment, and the generation of specialized pro-resolving mediators. Accordingly, a lower AA/DHA ratio indicates a relatively greater contribution of DHA to the long-chain PUFA pool, while a higher ratio indicates relative predominance of AA-dependent lipid signaling potential. This index is particularly informative in matrices where both fatty acids are present at nutritionally relevant levels, such as human milk, infant formulas, marine foods, DHA-enriched egg products, and biological lipidomes enriched in membrane phospholipids. However, the ratio should not be interpreted as simply "high = bad" and "low = good", because AA is also an essential fatty acid with important physiological roles, and the optimal balance depends on age, tissue, nutritional context, inflammatory status, and biological matrix. In NMLP, AA/DHA is used as an exploratory omega-6/omega-3 balance descriptor focused on the relative contribution of AA- versus DHA-related lipid biology. | 30, 31 |
| | EPA_DHA | EPA + DHA | EPA + DHA = C20:5n-3 + C22:6n-3 | An increased EPA + DHA value indicates greater representation of long-chain omega-3 fatty acids. | A decreased EPA + DHA value indicates lower representation of long-chain omega-3 fatty acids. | The EPA + DHA index, defined as the sum of eicosapentaenoic acid (20:5n-3) and docosahexaenoic acid (22:6n-3), expresses the total representation of the two principal bioactive long-chain omega-3 polyunsaturated fatty acids within a lipid matrix. It can be interpreted as a descriptor of long-chain omega-3 status. EPA and DHA are key constituents of cellular membranes and are directly linked to the structural, signaling, and pro-resolving roles of long-chain n-3 PUFA in human physiology. Because conversion from alpha-linolenic acid to EPA and DHA is limited in humans, this index may also reflect exposure to preformed long-chain omega-3 fatty acids, particularly from marine-derived foods or supplements. However, in NMLP, EPA + DHA is primarily used to characterize the biological representation of long-chain omega-3 fatty acids in the lipidome, rather than the nutritional quality of foods. A higher EPA + DHA value is generally compatible with a more omega-3-enriched lipid profile, whereas a lower value indicates poorer representation of bioactive long-chain omega-3s. Its interpretation should be refined together with AA/DHA, n-6/n-3 balance, total PUFA, lipid class distribution, and oxidative susceptibility. | 32 |
| | LA_ALA_Ratio | LA/ALA Ratio | LA/ALA = C18:2n-6 / C18:3n-3 | An increased LA/ALA ratio indicates greater linoleic-acid predominance relative to alpha-linolenic acid, shifting the essential fatty-acid precursor balance toward n-6. | A decreased LA/ALA ratio indicates greater alpha-linolenic-acid representation relative to linoleic acid, shifting the essential fatty-acid precursor balance toward n-3. | The LA/ALA ratio, defined as 18:2n-6 divided by 18:3n-3, expresses the balance between the two principals essential PUFA precursors: linoleic acid of the n-6 series and alpha-linolenic acid of the n-3 series. It is best interpreted as an essential fatty-acid precursor balance descriptor rather than as a direct marker of a specific clinical outcome. Its biochemical relevance lies in the fact that LA and ALA share desaturation and elongation pathways involved in the endogenous synthesis of long-chain PUFA, so a higher LA/ALA ratio reflects relative predominance of the n-6 precursor and a less favourable substrate balance for conversion of ALA toward EPA and DHA. Conversely, a lower LA/ALA ratio indicates a more n-3-precursor-oriented lipid profile. This interpretation is especially relevant in infant nutrition, where historical recommendations for term formulas placed the LA/ALA ratio between approximately 5:1 and 15:1, and lowering the ratio from 10:1 to 5:1 was shown to produce a modest increase in DHA status, although DHA levels still remained below those observed in breast-fed infants. In broader nutritional-metabolic contexts, a lower LA/ALA ratio may support greater ALA availability and, in some interventions, higher EPA status, but endogenous long-chain n-3 PUFA synthesis remains limited, particularly in adults. Therefore, changes in LA/ALA should not be assumed to translate into large increases in EPA + DHA status. In NMLP, LA/ALA is used as an exploratory descriptor of the balance between essential n-6 and n-3 PUFA precursors. | 33, 34, 35 |
| | HUFA_like_GPL | HUFA-like GPL | abundance of GPL species with DB ≥ 4 / total GPL | An increased HUFA-like GPL fraction suggests enrichment of highly unsaturated glycerophospholipids; it should be interpreted as a HUFA-like remodeling proxy, not as a direct omega-3 marker. | A decreased HUFA-like GPL fraction suggests lower representation of highly unsaturated glycerophospholipids. | The HUFA-like GPL index is defined as the abundance of glycerophospholipid species with DB ≥ 4 divided by total glycerophospholipids. It estimates how much of the GPL pool is represented by species carrying multiple double bonds, often compatible with long-chain PUFA-rich phospholipids. Glycerophospholipids are major membrane lipids, and their acyl-chain unsaturation strongly influences membrane packing, flexibility, curvature, and deformation behaviour. In biochemical and nutritional terms, this index can preserve some interpretability of highly unsaturated fatty-acid incorporation into glycerophospholipids when explicit fatty-acid resolution is not available. However, it should be read as a HUFA-like GPL proxy rather than as a direct omega-3 index, because it does not distinguish n-3 from n-6 highly unsaturated species. A higher HUFA-like GPL fraction is therefore consistent with a GPL pool relatively enriched in highly unsaturated phospholipids, which may reflect dietary unsaturated-fat exposure, PUFA incorporation into complex lipids, or membrane lipid remodelling. At the same time, highly unsaturated glycerophospholipids are more vulnerable to lipid peroxidation, so the biological meaning of this index remains context dependent. In NMLP, HUFA-like GPL is used as an exploratory structural | 36 |

| | | | | | | | |
|---|---|---|---|---|---|---|---|
| | | | | | | proxy of highly unsaturated GPL remodelling, to be interpreted together with EPA + DHA, AA/DHA, LA/ALA, n-6/n-3 balance, and oxidative-susceptibility indices. | |
| | Very_HUFA_GPL | Very_HUFA_GPL | GPL con DB >= 5 / GPL total | An increased Very-HUFA GPL fraction suggests enrichment of the most highly unsaturated glycerophospholipids. | A decreased Very-HUFA GPL fraction suggests lower representation of the most highly unsaturated glycerophospholipids. | The Very_HUFA_GPL index is defined as the abundance of glycerophospholipid species with DB ≥ 5 divided by total glycerophospholipids. It estimates the relative contribution of the most highly unsaturated GPL species within the membrane phospholipid pool. Higher values are generally compatible with greater representation of long-chain PUFA-rich membrane lipids and with a lipidome more oriented toward highly unsaturated remodelling. Glycerophospholipids are major membrane lipids, and high acyl-chain unsaturation strongly affects membrane flexibility, curvature responses, bilayer packing, and lipid-mediated signaling. This index can preserve some interpretability of very highly unsaturated PUFA incorporation into glycerophospholipids when explicit fatty-acid resolution is not available. However, DB ≥ 5 is an operational threshold rather than a direct biochemical assignment, and the index does not by itself distinguish omega-3 from omega-6 highly unsaturated chains. Very highly unsaturated glycerophospholipids are also more susceptible to lipid peroxidation, so the biological meaning of this index remains context dependent. In NMLP, Very_HUFA_GPL is used as an exploratory structural proxy of very highly unsaturated GPL remodelling, to be interpreted together with EPA + DHA, AA/DHA, LA/ALA, n-6/n-3 balance, and oxidative-susceptibility indices. | 36 |
| | Low_unsat_GPL | Low_unsat_GPL | GPL con DB <= 2 / GPL total | An increased Low-unsat GPL fraction suggests greater representation of low-unsaturation glycerophospholipids. | A decreased Low-unsat GPL fraction suggests lower representation of low-unsaturation glycerophospholipids and a relatively greater contribution of more unsaturated GPL species. | The Low_unsat_GPL index, defined as the abundance of glycerophospholipid species with DB ≤ 2 divided by total glycerophospholipids, can be interpreted as a proxy descriptor of low-unsaturation GPL abundance. Higher values are generally compatible with a membrane lipid pool less enriched in highly unsaturated species and more weighted toward saturated, monounsaturated, or weakly unsaturated structural components. This interpretation is biochemically plausible because glycerophospholipid unsaturation strongly influences membrane packing, flexibility, curvature, and deformation behaviour. Compared with HUFA-like GPL-rich profiles, a higher Low_unsat_GPL fraction is generally more compatible with tighter acyl-chain packing and lower contribution of long-chain PUFA-rich species to membrane architecture. This index can preserve some interpretability of membrane unsaturation balance when explicit fatty-acid resolution is not available. However, DB ≤ 2 is an operational threshold rather than a direct biochemical assignment, and the index does not distinguish specific fatty-acid species or n-3/n-6 origin. Directionality is descriptive rather than intrinsically favourable or unfavourable: higher values may indicate lower PUFA/HUFA representation but also lower oxidative susceptibility, whereas lower values may indicate greater unsaturated GPL remodelling but potentially higher peroxidability. In NMLP, Low_unsat_GPL is used as a complementary proxy of low-unsaturation GPL representation, to be interpreted together with HUFA-like GPL, Very-HUFA GPL, EPA + DHA, AA/DHA, n-6/n-3 balance, and oxidative-susceptibility indices. | 36 |
| | HUFA_shift_GPL | HUFA_shift_GPL | (GPL con DB >= 4) / (GPL con DB <= 2) | An increased HUFA-shift GPL ratio suggests a displacement of the GPL pool toward highly unsaturated species relative to low-unsaturation species. | A decreased HUFA-shift GPL ratio suggests a displacement of the GPL pool toward lower-unsaturation species. | The HUFA_shift_GPL index, defined as the abundance of glycerophospholipid species with DB ≥ 4 divided by the abundance of glycerophospholipid species with DB ≤ 2, can be interpreted as a proxy descriptor of the relative shift from low unsaturation to highly unsaturated GPL species. In practical terms, it provides a synthetic view of whether the membrane GPL pool is displaced toward a more HUFA-like unsaturation pattern or toward a lower-unsaturation structural pattern. This interpretation is biochemically plausible because glycerophospholipid unsaturation strongly influences membrane packing, flexibility, curvature, deformation behaviour, and lipid-mediated signaling. A higher HUFA_shift_GPL ratio is generally compatible with a GPL pool richer in highly unsaturated species relative to low-unsaturation ones, whereas a lower ratio is compatible with a GPL pool more weighted toward saturated, monounsaturated, or weakly unsaturated structural components. However, DB ≥ 4 and DB ≤ 2 are operational thresholds rather than direct biochemical assignments, and this index does not distinguish omega-3 from omega-6 highly unsaturated species. It should therefore be interpreted as a remodelling proxy, not as a direct omega-3 marker. Its biological meaning remains context-dependent because highly unsaturated GPLs may support membrane adaptability and PUFA-related signaling, but they may also increase susceptibility to lipid peroxidation. The ratio should also be handled carefully when the low-unsaturation GPL denominator is very small. In NMLP, HUFA_shift_GPL is used as an exploratory descriptor of the balance between highly unsaturated and low-unsaturation GPL species. | 36 |
| Overall Lipid Quality | PUFA_SFA_Ratio | PUFA/SFA Ratio | PUFA/SFA = ΣPUFA / ΣSFA | An increased PUFA/SFA ratio suggests a lipid profile relatively enriched in polyunsaturated compared with saturated fatty acids, generally compatible with higher unsaturation-based lipid quality. | A decreased PUFA/SFA ratio suggests a lipid profile relatively enriched in saturated compared with polyunsaturated fatty acids, generally compatible with lower unsaturation-based lipid quality. | The PUFA/SFA ratio, defined as total polyunsaturated fatty acids divided by total saturated fatty acids, expresses the balance between two major fatty acid groups relevant to lipid metabolism, membrane composition, dietary fat quality, and cardiovascular health. A higher value indicates a lipid fraction relatively enriched in PUFA and less dominated by SFA, corresponding to a more unsaturated lipid profile with potential implications for membrane properties, lipid mediator availability, and overall lipid quality. This ratio is one of the most widely used indices for evaluating fat quality, especially in food profiling and dietary intervention studies, where higher PUFA/SFA values are generally interpreted as more favourable for cardiovascular health, particularly when accompanied by meaningful n-3 PUFA, EPA, and DHA contributions. However, the ratio should not be interpreted in isolation: its biological meaning depends on the n-6/n-3 balance, oxidative susceptibility, lipid class distribution, biological matrix, and metabolic context. In NMLP, PUFA/SFA is therefore used as a broad unsaturation-based lipid quality descriptor rather than as a standalone clinical or nutritional score. | 37, 38 |
| | SFA_Fraction | SFA Fraction | SFA/frac = ΣSFA / Σtotal fatty acids | An increased SFA fraction indicates a greater relative contribution of saturated fatty acids, generally suggesting a less favorable unsaturation-based lipid quality profile. | A decreased SFA fraction indicates a lower relative contribution of saturated fatty acids, generally compatible with greater replacement by unsaturated fatty acids. | The SFA Fraction index, defined as total saturated fatty acids divided by total fatty acids, reflects the relative contribution of saturated fatty acids within the lipid compartment under study. It is best interpreted as a broad saturated-fat enrichment descriptor with relevance to overall lipid quality and cardiometabolic interpretation, rather than as a standalone cardiovascular risk marker. A higher value indicates a lipid pool more dominated by saturated fatty acids and is generally considered less favourable when it reflects insufficient replacement of SFA with unsaturated fatty acids. Conversely, a lower SFA fraction is usually interpreted as more favourable when it reflects substitution of SFA with MUFA or PUFA. However, the biological meaning of this index depends on the lipid matrix, dietary context, replacement nutrient, and fatty-acid sources. In NMLP, SFA Fraction is therefore used as a general descriptor of saturated-fat contribution within the lipidome, complementary to PUFA/SFA, MUFA/SFA, and other unsaturation-based quality indices. | 39, 40 |
| | ACL_sum | ACL/sum | Σ(abundance × total carbons) / Σ(abundance) | Higher ACL/sum suggests a lipid pool composed of species with higher average total carbon | Lower ACL/sum suggests a lipid pool composed of species with lower average | The ACL/sum index can be interpreted as the abundance-weighted average total carbon content per lipid species, calculated at the sum-composition level. It describes the average total number of carbons of the annotated lipid species in the selected lipid set and therefore functions as a proxy for the overall molecular carbon size of the lipid pool. A higher ACL/sum indicates that the lipid pool is, on average, composed of species with more total | 41 |

| | | | | | | | |
|---|---|---|---|---|---|---|---|
| | | | | number, within the limits of SUM-level annotation. | total carbon number, within the limits of SUM-level annotation. | carbons, whereas a lower ACL/sum indicates species with fewer total carbons. ACL/sum should be regarded as a molecule-level structural descriptor of lipid composition rather than as a direct measure of the average length of each acyl chain. This distinction is important because the classical ACL/chain metric explicitly normalizes by the number of fatty acyl chains, whereas ACL/sum does not. For this reason, ACL/sum is useful as a descriptive index of the average size of annotated lipid species, particularly in SUM-level datasets, but it is less immediately interpretable biologically than ACL/chain when lipid classes with different molecular architectures are analysed together. Its interpretation should therefore remain cautious and context-dependent, because ACL/sum is influenced by chain length, number of acyl chains per lipid class, lipid-class composition, and the structural resolution of the annotation. | |
| | DB_C_Sum | DB/C Sum | total double bonds / total carbons | Higher DB/C Sum suggests greater unsaturation density relative to total carbon content in the lipid pool. | Lower DB/C Sum suggests lower unsaturation density relative to total carbon content in the lipid pool. | DB/C Sum, calculated as the abundance-weighted ratio of total double bonds to total carbon atoms across the lipid pool, expresses the average unsaturation density of the lipidome. It captures whether the overall lipid composition is shifted toward relatively more saturated or more unsaturated molecular species after normalization for total carbon content. A higher DB/C Sum indicates a greater number of double bonds per carbon and is compatible with enrichment in MUFA- and PUFA-containing lipids, generally reflecting a more unsaturated lipid-quality profile. Conversely, a lower DB/C Sum reflects lower average unsaturation density, consistent with enrichment in more saturated or less unsaturated lipid species. This index can therefore be interpreted as a global lipid-quality balance descriptor, while also providing information potentially relevant to membrane behaviour, lipid packing, and remodelling. However, DB/C Sum should not be interpreted as uniformly favourable or unfavourable, because greater unsaturation may also increase chemical susceptibility to oxidation, and its biological meaning depends on the lipid classes involved, the n-3/n-6 balance, the biological matrix, and the oxidative/metabolic context. In NMLP, DB/C Sum is used as a broad descriptor of whole-lipidome unsaturation density. | 42, 43, 44 |
| | DFA | Desirable Fatty Acids (DFA) | MUFA + PUFA + C18:0 | Higher DFA indicates a greater contribution of fatty acids generally considered desirable for fat-quality assessment. | Lower DFA indicates a smaller contribution of fatty acids generally considered desirable for fat-quality assessment. | The Desirable Fatty Acids index represents the sum of fatty acids generally considered nutritionally more favourable for fat-quality assessment, conventionally calculated as MUFA + PUFA + C18:0. Its rationale is based on the inclusion of unsaturated fatty acids, which are generally considered more favourable than the major hypercholesterolemic saturated fatty acids, together with stearic acid, a saturated fatty acid with a substantially more neutral effect on LDL cholesterol than lauric, myristic, and palmitic acids. A higher DFA value indicates that a greater proportion of the fat fraction is composed of fatty acids regarded as desirable from a nutritional perspective, whereas a lower DFA value reflects a less favourable fatty-acid composition. DFA is commonly interpreted together with the complementary OFA index or the DFA/OFA ratio. It is used primarily as a food lipid-quality index, especially in studies on oils, meat, milk, fish, and infant formulas, where it serves to characterize the compositional balance between desirable and hypercholesterolemic fatty acids rather than to estimate a specific biological pathway or clinical outcome directly. In NMLP, DFA should therefore be interpreted as a broad fat-quality descriptor, with its meaning refined by complementary indices such as PUFA/SFA, n-6/n-3, EPA+DHA, SFA Fraction, and oxidative susceptibility indices. | 42, 44, 45 |
| | MUFA_Fraction | MUFA Fraction | MUFA/frac = ΣMUFA / Σtotal fatty acids | An increased MUFA fraction indicates a greater relative contribution of monounsaturated fatty acids to the total fatty-acid pool. | A decreased MUFA fraction indicates a lower relative contribution of monounsaturated fatty acids to the total fatty-acid pool. | The MUFA Fraction index, commonly expressed as ΣMUFA / Σtotal fatty acids, quantifies the relative contribution of monounsaturated fatty acids to the total fatty-acid pool. It is best interpreted as a compositional descriptor of lipid quality balance, because it describes how strongly a lipid profile is shifted toward MUFA-rich composition rather than toward saturated or polyunsaturated fatty acids. Biochemically, however, the meaning of a higher MUFA fraction is context-dependent, since an elevated MUFA contribution may reflect either greater dietary exposure to MUFA-rich foods or enhanced endogenous desaturation and lipogenesis, particularly through pathways involving stearoyl-CoA desaturase activity in metabolically altered tissues. When an increase in MUFA Fraction accompanies the replacement of saturated fatty acids with unsaturated fatty acids, it is generally interpreted as favourable, in line with dietary intervention studies showing that improved dietary fat quality is associated with beneficial remodelling of the lipidome and cardiometabolic markers. In contrast, in internal biological compartments such as liver or other tissues affected by steatosis or metabolic dysfunction, a relatively high MUFA fraction may instead reflect pathological endogenous MUFA production rather than a beneficial dietary pattern. Accordingly, MUFA Fraction is most appropriately viewed as a compositional lipid-balance descriptor whose interpretation depends on whether the observed MUFA enrichment arises predominantly from dietary fat quality or from metabolic remodelling. | 16, 42 |
| | PUFA_Fraction | PUFA Fraction | PUFA/frac = ΣPUFA / Σtotal fatty acids | An increased PUFA fraction indicates a greater relative contribution of polyunsaturated fatty acids to the total fatty-acid pool. | A decreased PUFA fraction indicates a lower relative contribution of polyunsaturated fatty acids to the total fatty-acid pool. | The PUFA Fraction index, defined as total polyunsaturated fatty acids divided by total fatty acids, reflects the relative contribution of PUFA within the lipid compartment under study and can be interpreted as a broad descriptor of unsaturation-based lipid quality. A higher value indicates a lipid pool relatively enriched in polyunsaturated fatty acids, a compositional feature relevant for membrane properties, lipid packing, and the availability of precursors for eicosanoid- and docosanoid-related pathways. This index is commonly used to describe how strongly a matrix is enriched in PUFA relative to the total fatty-acid pool, and higher PUFA proportions are generally interpreted as nutritionally favourable, especially when accompanied by substantial n-3 PUFA, EPA, and DHA contributions. In dietary intervention and lipidomics studies, increasing dietary PUFA or replacing SFA with unsaturated fats can shift the plasma lipidome toward a more unsaturated profile, with remodelling particularly evident in glycerophospholipids, including ethanolamine-containing and plasmalogen species. However, PUFA Fraction should not be interpreted in isolation, because its biological meaning depends on the n-6/n-3 balance, EPA/DHA contribution, lipid class distribution, biological matrix, and oxidative environment. Greater PUFA enrichment may improve lipid quality and membrane adaptability, but it can also increase susceptibility to lipid peroxidation. In NMLP, PUFA Fraction is therefore used as a broad lipid-quality descriptor of polyunsaturated fatty-acid contribution. | 42, 46, 47 |
| | High_DB_fraction | High_DB_fraction | abundance of species with DB ≥ 4 / total abundance | An increased High-DB fraction indicates greater representation of highly unsaturated lipid species, compatible with greater lipid unsaturation and potentially higher oxidative susceptibility. | A decreased High-DB fraction indicates lower representation of highly unsaturated lipid species. | The High_DB_fraction index, defined as the abundance of lipid species with DB ≥ 4 divided by total lipid abundance, can be interpreted as a global descriptor of highly unsaturated lipid species within the lipidome. It estimates how much of the lipid pool is represented by lipids carrying multiple double bonds, often including PUFA-rich molecular species. These highly unsaturated lipids are biologically important because they can contribute to membrane adaptability, bilayer flexibility, deformation capacity, and lipid-mediated signaling. A higher High_DB_fraction may also be compatible with greater representation of diet-related unsaturated fats, especially long-chain n-3 and n-6 PUFA-containing lipids. At the same time, highly unsaturated lipids are more susceptible to lipid peroxidation and oxidative remodelling. For this reason, High_DB_fraction should be | 12, 36 |

| | | | | | | | |
|---|---|---|---|---|---|---|---|
| | | | | | | interpreted as a lipid quality and remodelling descriptor, not as a uniformly favourable or unfavourable index. Its biological meaning depends on lipid class distribution, PUFA type, oxidative environment, biological matrix, and annotation resolution. In NMLP, High_DB_fraction is therefore used as an exploratory descriptor of the relative contribution of highly unsaturated lipid species to the overall lipidome. | |
| | UI | Unsaturation Index (UI) | UI = 1×(% monoenoics) + 2×(% dienoics) + 3×(% trienoics) + 4×(% tetraenoics) + 5×(% pentaenoics) + 6×(% hexaenoics) | An increased UI indicates a higher overall degree of fatty-acid unsaturation, generally reflecting a more unsaturated lipid profile. | A decreased UI indicates a lower overall degree of fatty-acid unsaturation. | The Unsaturation Index is a weighted measure of the overall degree of fatty-acid unsaturation in a lipid mixture, calculated by multiplying the proportion of each unsaturated fatty-acid group by its number of double bonds. In this way, pentaenoic and hexaenoic fatty acids contribute more strongly than monoenoic or dienoic fatty acids, allowing UI to summarize not only how many unsaturated fatty acids are present, but also how highly unsaturated the lipid pool is. UI is therefore a compact structural descriptor of food lipids, cellular membranes, tissue lipid stores, or plasma lipidomes. Biologically, a higher UI indicates a profile enriched in more highly unsaturated fatty acids and is generally compatible with greater membrane adaptability, conformational flexibility, altered lipid packing, and increased availability of PUFA-derived bioactive precursors. Conversely, a lower UI reflects a relatively less unsaturated and structurally more saturated lipid environment. Because susceptibility to lipid peroxidation increases with the number of double bonds, UI also has secondary relevance for oxidative susceptibility; however, it should be interpreted primarily as an unsaturation descriptor rather than as a specific oxidation index. Its biological meaning remains context-dependent and should be evaluated together with PUFA type, n-6/n-3 balance, EPA/DHA contribution, lipid class distribution, biological matrix, and oxidative environment. In NMLP, UI is used as a broad lipid-quality descriptor of the overall unsaturation state of the lipid pool. | 48, 49 |
| Oxidative Protection | PC_PE_Ratio | PC/PE Ratio | PC/PE = Σ phosphatidylcholine / Σ phosphatidylethanolamine | An increased PC/PE ratio suggests a shift in membrane phospholipid balance toward phosphatidylcholine; interpretation is compartment- and context-dependent. | A decreased PC/PE ratio suggests a shift in membrane phospholipid balance toward phosphatidylethanolamine; interpretation is compartment- and context-dependent. | The PC/PE ratio, expressed as total phosphatidylcholine divided by total phosphatidylethanolamine, quantifies the balance between two major structural phospholipid classes in biological membranes. It is best interpreted as a membrane phospholipid balance and membrane-integrity descriptor rather than as a classical dietary fat-quality index or a direct oxidative-protection marker. PC and PE have distinct biophysical roles: PC favours a more lamellar, bilayer-stabilizing geometry, whereas PE, owing to its smaller headgroup, promotes curvature stress and different packing properties within membranes. As a result, the PC/PE ratio is an important determinant of membrane organization, permeability, and structural adequacy. In liver, this ratio is closely linked to PC synthesis from dietary choline through the CDP-choline pathway and to PE methylation through PEMT, which converts PE into PC. When choline availability is inadequate or PC synthesis is impaired, the PC/PE ratio can decrease, compromising membrane integrity and hepatic function. A low PC/PE ratio has been associated with impaired membrane integrity, increased permeability, and progression from steatosis to steatohepatitis/NASH, especially in hepatic settings. A higher ratio may be beneficial when it reflects restoration from an abnormally low baseline, such as under choline deficiency or hepatic phosphatidylcholine insufficiency. However, the interpretation is not universally linear: in some metabolic contexts linked to obesity, insulin resistance, or type 2 diabetes, a higher PC/PE ratio may coexist with other unfavourable lipidomic signals. Because PC/PE balance is tissue-, compartment-, and disease-dependent, this index should be interpreted as a membrane-balance descriptor with indirect relevance to stress and oxidative vulnerability, rather than as an invariably favourable or unfavourable marker. | 50, 51, 52 |
| | Ether_PC_Total_PC | Ether-PC / Total PC | (PC-O + PC-P) / total PC | An increased Ether-PC/Total PC ratio suggests enrichment of ether-linked and plasmalogen PC species within the total PC pool. | A decreased Ether-PC/Total PC ratio suggests relative depletion of ether-linked and plasmalogen PC species within the total PC pool. | The Ether-PC/Total PC index, defined as the sum of PC-O and PC-P divided by total PC, reflects the relative contribution of ether-linked and plasmalogen phosphatidylcholines within the total phosphatidylcholine pool. It is best interpreted as a marker of the ether/plasmalogen signature of the PC class rather than as a general measure of total PC abundance. A higher value indicates that the PC pool is relatively enriched in alkyl- and alkenyl-ether species, which are structurally and functionally distinct from conventional diacyl-PC and have been linked to membrane organization, lipid packing, lateral membrane behaviour, peroxisomal lipid metabolism, and redox-related functions. In food composition studies, this index can describe how strongly an animal-derived matrix is enriched in ether-PC or plasmalogen-PC species, consistent with reports showing that choline plasmalogens occur in meats, poultry, seafood, and other animal-derived products. In biological samples, the same ratio can capture diet- or metabolism-related remodelling of the ether-PC branch. A higher Ether-PC/Total PC ratio may therefore indicate relative enrichment of the PC pool in species potentially linked to membrane adaptation and ether/plasmalogen-associated redox resilience, whereas a lower ratio suggests depletion of the ether-linked PC compartment and potentially lower plasmalogen-associated protection. However, the interpretation remains context-dependent because PC-O and PC-P are not equivalent, and the biological meaning depends on lipid class distribution, sample type, diet, peroxisomal metabolism, oxidative environment, and annotation quality. In NMLP, Ether-PC/Total PC is used as an exploratory oxidative-protection and membrane-remodelling descriptor of the ether-linked PC fraction. | 53 |
| | Ether_PE_Total_PE | Ether-PE / Total PE | (PE-O + PE-P) / total PE | An increased Ether-PE/Total PE ratio suggests enrichment of ether-linked and plasmalogen PE species within the total PE pool. | A decreased Ether-PE/Total PE ratio suggests relative depletion of ether-linked and plasmalogen PE species within the total PE pool. | The Ether-PE/Total PE index, defined as the sum of PE-O and PE-P divided by total PE, reflects the relative contribution of alkyl-ether and plasmalogen phosphatidylethanolamines within the total phosphatidylethanolamine pool. It is best interpreted as a marker of the ethanolamine ether/plasmalogen signature of the PE class rather than as a general measure of total PE abundance. A higher value indicates that the PE pool is relatively enriched in ether-linked and vinyl-ether species, which are biophysically distinct from conventional diacyl-PE and have been linked to membrane curvature, fusion–fission dynamics, lipid domain organization, peroxisomal lipid metabolism, and redox-related membrane adaptation. In food composition studies, this index can describe how strongly an animal-derived matrix is enriched in ether-PE/plasmalogen-PE, consistent with evidence that dietary plasmalogens occur in meat, poultry, fish, molluscs, and other phospholipid-rich animal foods. In biological samples, the same ratio can capture diet- and metabolism-sensitive remodelling of the ether-PE branch. A higher Ether-PE/Total PE ratio may therefore indicate relative enrichment of the PE pool in species potentially linked to membrane specialization and ether/plasmalogen-associated redox resilience, whereas a lower ratio suggests depletion of the ether-linked PE compartment and potentially lower plasmalogen-associated membrane protection. However, the interpretation remains context-dependent because PE-O and PE-P are not equivalent, and the biological meaning depends on lipid class distribution, sample type, diet, peroxisomal metabolism, oxidative environment, and annotation quality. In NMLP, Ether-PE/Total PE is used as an exploratory oxidative-protection and membrane-remodelling descriptor of the ether-linked PE fraction. | 53, 54 |

| | | | | | | | |
|---|---|---|---|---|---|---|---|
| | PS_Total_GPL | PS / Total GPL | PS/GPL = total phosphatidylserine / total glycerophospholipids | An increased PS/Total GPL ratio indicates greater relative contribution of phosphatidylserine within the glycerophospholipid pool. | A decreased PS/Total GPL ratio indicates lower relative contribution of phosphatidylserine within the glycerophospholipid pool. | The PS/GPL ratio, defined as total phosphatidylserine divided by total glycerophospholipids, reflects the relative contribution of phosphatidylserine within the overall glycerophospholipid pool. It is best interpreted as a membrane phospholipid balance and anionic phospholipid signaling descriptor rather than as a direct oxidative-protection marker. PS is a negatively charged glycerophospholipid synthesized in the endoplasmic reticulum and mitochondria-associated membranes from major GPL classes such as PC and PE through the action of PSS1 and PSS2. It is enriched in the cytosolic leaflet of the plasma membrane and in selected intracellular compartments, where it contributes to membrane asymmetry, apoptotic and efferocytic signaling, procoagulant membrane activity, cholesterol trafficking, and surface-dependent signaling processes. A higher PS/GPL ratio indicates a relatively greater representation of PS within total GPL, consistent with a more PS-enriched membrane profile. In some contexts, such as metabolic liver disease, hepatic PS accumulation and preserved PSS1 activity have been linked to VLDL remodelling, dyslipidemia, and hypertriglyceridemia, suggesting that increased relative PS may be metabolically unfavourable in specific settings. Conversely, a lower PS/GPL ratio suggests a smaller relative PS contribution to total GPL, but this should not be interpreted as automatically beneficial because PS remains essential for membrane organization and signaling. Because PS changes may reflect membrane composition, signaling, vesicle biology, apoptosis-related processes, or metabolic remodelling, the directionality of this index should be interpreted in the biological context of the study. | 55 |
| | PEP_PE_Ratio | PEP/PE Ratio | PEP/PE = Σ plasmalogen PE / Σ total PE | An increased PEP/PE ratio indicates greater relative abundance of PE plasmalogens within the PE pool, generally compatible with stronger plasmalogen representation. | A decreased PEP/PE ratio indicates relative depletion of PE plasmalogens within the PE pool. | The PEP/PE ratio, defined as total plasmalogen phosphatidylethanolamine [PE(P)] divided by total phosphatidylethanolamine [PE], reflects the relative contribution of ethanolamine plasmalogens within the overall PE pool. It can therefore be interpreted as a marker of how strongly the phosphatidylethanolamine compartment is shifted toward a plasmalogen-rich phenotype. Biochemically, this ratio is informative because PE plasmalogens are ether phospholipids characterized by a vinyl-ether bond that confers distinctive properties related to membrane organization, lipid packing, membrane fluidity, and redox biology. From a nutritional and metabolic perspective, recent human lipidomics evidence supports the relevance of PE(P) relative to PE: the plasmalogen score developed by Beyene et al. captures a metabolic signal based on PE(P) and PE, shows its strongest correlation with the PE(P)/PE ratio, is inversely associated with type 2 diabetes, cardiovascular disease, and all-cause mortality, and is modifiable by diet and lifestyle. Accordingly, a higher PEP/PE ratio is generally consistent with a more favourable plasmalogen-rich PE phenotype and potentially greater membrane/redox resilience, whereas a lower ratio suggests relative plasmalogen depletion within the PE pool, a pattern compatible with poorer cardiometabolic health in circulating lipidomes. However, this index should still be interpreted cautiously because its meaning depends on biological matrix, diet, lifestyle, peroxisomal metabolism, oxidative environment, disease context, and annotation quality. In NMLP, PEP/PE is used as an exploratory oxidative-protection and cardiometabolic-resilience descriptor of PE plasmalogen representation. | 54 |
| | SM_Cer_Ratio | SM/Cer Ratio | SM/Cer = Σ sphingomyelin / Σ ceramide | An increased SM/Cer ratio suggests sphingomyelin predominance relative to ceramide, generally compatible with a lower relative ceramide burden. | A decreased SM/Cer ratio suggests relative ceramide enrichment, generally compatible with a less favourable sphingolipid balance. | The SM/Cer ratio, defined as total sphingomyelin divided by total ceramide within the same biological or food compartment, reflects the balance between sphingomyelin, a largely structural sphingolipid of membranes and lipoproteins, and ceramide, a more bioactive lipid class linked to lipotoxic, inflammatory, and cardiometabolic signaling. It is therefore best interpreted as an exploratory sphingolipid-balance descriptor with cardiometabolic risk relevance, rather than as a direct clinical risk score. Biochemically, this ratio is meaningful because ceramide can arise both from de novo synthesis and from sphingomyelin hydrolysis by sphingomyelinases. From a nutritional and metabolic perspective, a higher SM/Cer ratio can indicate a profile relatively shifted toward the structural sphingomyelin compartment and away from ceramide accumulation, whereas a lower ratio is more consistent with ceramide enrichment and a less favourable sphingolipid balance. This interpretation is supported by the broader literature linking elevated ceramides to cardiovascular disease, obesity, insulin resistance, and diabetes. Diet can also modulate circulating and lipoprotein sphingolipids: saturated-fat-rich exposures, particularly palmitate-related pathways, have been associated with increased ceramide production, whereas dietary sphingolipids and milk polar lipids may modify sphingomyelin and ceramide profiles. However, the ratio should remain context-dependent, because sphingomyelin and ceramide species differ by acyl-chain composition, biological compartment, lipoprotein distribution, and disease context. In NMLP, SM/Cer is used as an exploratory descriptor of sphingomyelin-to-ceramide balance and relative ceramide burden. | 56, 57, 58 |

(55) Anari, M.; Karimkhanloo, H.; Nie, S.; Dong, L.; Fidelito, G.; Bayliss, J.; Keenan, S. N.; Slavin, J.; Lin, S.; Cheng, Z.; Lu, J.; Miotto, P. M.; De Nardo, W.; Devereux, C. J.; Williamson, N. A.; Watt, M. J.; Montgomery, M. K. Lipidome Profiling in Advanced Metabolic Liver Disease

Identifies Phosphatidylserine Synthase 1 as a Regulator of Hepatic Lipoprotein Metabolism. *Cell Reports* **2024**, *43* (12), 115007. https://doi.org/10.1016/j.celrep.2024.115007.

(56) Boulgaropoulos, B.; Amenitsch, H.; Laggner, P.; Pabst, G. Implication of Sphingomyelin/Ceramide Molar Ratio on the Biological Activity of Sphingomyelinase. *Biophysical Journal* **2010**, *99* (2), 499–506. https://doi.org/10.1016/j.bpj.2010.04.028.

(57) Wang, S.; Jin, Z.; Wu, B.; Morris, A. J.; Deng, P. Role of Dietary and Nutritional Interventions in Ceramide-Associated Diseases. *Journal of Lipid Research* **2025**, *66* (1), 100726. https://doi.org/10.1016/j.jlr.2024.100726.

(58) Voß, J.; Hornemann, T.; Belgardt, B. Impact of Nutrition on Sphingolipid-Regulated Physiology: A Review. *Molecular Nutrition Food Res* **2025**, *69* (20), e70182. https://doi.org/10.1002/mnfr.70182.

(59) Hachem, M.; Nacir, H. Emerging Role of Phospholipids and Lysophospholipids for Improving Brain Docosahexaenoic Acid as Potential Preventive and Therapeutic Strategies for Neurological Diseases. *IJMS* **2022**, *23* (7), 3969. https://doi.org/10.3390/ijms23073969.
